# Hypertrophic cardiomyopathy mutations Y115H and E497D disrupt the folded-back state of human β-cardiac myosin allosterically

**DOI:** 10.1101/2024.02.29.582851

**Authors:** Neha Nandwani, Debanjan Bhowmik, Matthew Carter Childers, Rama Reddy Goluguri, Aminah Dawood, Michael Regnier, James A. Spudich, Kathleen M. Ruppel

## Abstract

At the molecular level, clinical hypercontractility associated with many hypertrophic cardiomyopathy (HCM)-causing mutations in β-cardiac myosin appears to be driven by their disruptive effect on the energy-conserving, folded-back, super relaxed (SRX) ‘OFF’-state of myosin. A pathological increase in force production results from release of heads from this ‘OFF’-state, which results in an increase in the number of heads free to interact with actin and produce force. Pathogenic mutations in myosin can conceivably disrupt the ‘OFF’-state by (1) directly affecting the intramolecular interfaces stabilizing the folded-back state, or (2) allosterically destabilizing the folded-back state *via* disruption of diverse conformational states of the myosin motor along its chemomechanical cycle. However, very little is understood about the mutations that fall in the latter group. Here, using recombinant human β-cardiac myosin, we analysed the biomechanical properties of two such HCM-causing mutations, Y115H (in the transducer) and E497D (in the relay helix), neither of which falls in the regions that interact to stabilize myosin’s folded-back state. We find these mutations have diverse effects on the contractility parameters of myosin, yet the primary hypercontractile change in both cases is the destabilization of the ‘OFF’-state of myosin and increased availability of active myosin heads for actin-binding. Experimental data and molecular dynamics simulations indicate that these mutations likely destabilize the pre-powerstroke state of myosin, the conformation the motor adopts in the inactive folded-back state. We propose that destabilization of the folded-back state of myosin, directly and/or allosterically, is the molecular basis of hypercontractility in HCM in a far greater number of pathogenic mutations than currently thought.

## INTRODUCTION

Hypertrophic cardiomyopathy (HCM) is a heart muscle disease that affects more than 1 in 500 individuals and is associated with significant morbidity in the form of arrythmia, heart failure and sudden death^1–3^. It is clinically characterized by diastolic abnormalities and hypertrophic remodelling of the left ventricle. HCM is caused overwhelmingly by autosomal dominant mutations in sarcomeric genes, most commonly *MYH7* (encoding the heavy chain of β-cardiac myosin, the predominant ventricular motor) and *MYBPC3* (encoding cardiac myosin binding protein-C, cMyBPC)^4–6^. It is not completely clear how these point mutations in myosin and other proteins lead to hyperdynamic contraction of the heart and increased energy consumption^7,8^.

Muscle contraction is fuelled by ATP-dependent cyclic interactions between myosin-based thick filaments and actin-based thin filaments in sarcomeres. Myosin molecules that are not bound to actin filaments exist structurally in at least two distinct states: an ‘ON’ state in which heads are away from the thick filament backbone and available to interact with actin, and a folded-back ‘OFF’-state known as the interacting head motif (IHM) where the heads are not available for interaction with actin^9–11^. These structural states are thought to correlate with two distinct enzymatic states of myosin: a disordered relaxed state (DRX) with heads that hydrolyse ATP at the normal basal ATPase rate (which is ~100-fold slower than the actin-activated ATPase rate), and an energy conserving state called the super-relaxed state (SRX) with heads turning over ATP ~10-fold slower than DRX myosin^12–14^.

Myosin autoinhibition has been proposed to be an energy conserving mechanism by decreasing ATP consumption when myosin molecules are in their IHM state^14–16^. It is thought that for many myosin molecules the SRX to DRX conversion occurs with each beat of the heart, with the heads in the IHM OFF-state during diastole and in the ON-state during systole^17^. In cardiac muscle, however, ~50% of all myosin heads adopt the SRX conformation even in actively contracting sarcomeres^15,16^, and this pool might be dynamically regulated by several physiological stimuli, including regulatory light chain phosphorylation ^13,16,18–25^. The conversion between the DRX and SRX-IHM states is therefore expected to be a key determinant of cardiac contractility and energy utilization^15^. Not surprisingly then, perturbation of this conversion is linked to cardiovascular diseases^7,22,26^. Indeed, many HCM-causing mutations in β-cardiac myosin have been shown experimentally to result in hypercontractility by destabilizing the SRX-IHM ‘OFF’-state, which results in more myosin heads participating in systolic force production^6,8^.

In the IHM ‘OFF’-state, the two catalytic Subfragment 1 (S1) heads of the myosin dimer interact with each other and are folded back onto their proximal coiled-coil Subfragment 2 (S2) tail, which inhibits both ATP hydrolysis and actin-binding^10,11,27,28^. Over 700 HCM mutations throughout the sequence of *MYH7* have been reported^29^. Many of these are found on the surface of the interacting domains of the S1 heads, which includes a relatively flat surface termed the mesa^30^, and the proximal S2 tail of myosin^11^. Mapping these mutations on homology model(s) of the ‘OFF’ state of human cardiac myosin had previously suggested that many of them might destabilize the ‘OFF’ state by disrupting potential intramolecular head-head and head-tail ionic interactions sequestering the myosin heads, as well as intermolecular interactions with other sarcomeric proteins like cMyBPC that may further stabilize the ‘OFF’ state in the thick filaments^11,24,31,32^. Using recombinant human β-cardiac myosin protein and/or intact myofibers, several such myosin mutations have indeed been shown to increase the number of heads available for actin-binding using single-nucleotide turnover assays, actin-activated ATPase rate measurements, and fiber-diffraction studies^23,24,33–38^. This prediction is further supported by the recent high resolution (3.6 Å) cryo-EM structure of human β-cardiac myosin IHM, which provides near atomic-resolution details of the interactions stabilizing the IHM structure, allowing an understanding of how different HCM mutations might weaken it^39^.

It is immediately clear from the cardiac IHM structure that many HCM mutations are not directly located at the intramolecular interfaces stabilizing this sequestered state. How do such mutations lead to hypercontractility? While altered myosin chemo-mechanoenzyme activity might be one mechanism, it is unlikely to be the unifying theme across different mutations, as previous studies with recombinant mutant forms of human β-cardiac myosin have shown that HCM mutations do not consistently lead to hypercontractile changes in the fundamental biomechanical properties and kinetics of actively cycling myosin^6,8,26,40–43^. We therefore hypothesized that such HCM mutations can instead affect the stability of the sequestered IHM state allosterically. The myosin motor domain is highly allosteric^44^ and HCM mutations could affect the allosteric communication networks between different subdomains of the motor that lead to small conformational changes in the S1 heads, which results in weakening of the IHM structure. Predictions from a recent in silico study lend support to this hypothesis^31^, but there is little biophysical data exploring the effects of such mutations on the motor function of myosin and its ability to form the folded-back state.

Here, we investigate two previously uncharacterized pathogenic myosin HCM mutations, Y115H and E497D, neither of which lie on the intramolecular interfaces stabilizing the cardiac IHM structure. Tyr-115 lies on the first β-strand of the transducer, close to the active site, and Glu-497 lies in the relay helix (Fig. 1). Using human β-cardiac myosin protein, recombinantly expressed and purified, we find that as expected, mutations in these critical regions of the motor domain resulted in significant changes in the fundamental contractility parameters of myosin, although the two mutants were altered in their contractility parameters in drastically different ways. Remarkably, both Y115H and E497D mutations were found to severely compromise myosin’s ability to form the IHM ‘OFF’-state and increase the number of myosin heads functionally accessible for actin-binding. These results show that the disruption of the IHM ‘OFF’-state of myosin by HCM mutations is not limited to mutations in residues that are directly involved in stabilizing the IHM interfaces, in keeping with our unifying hypothesis that most myosin HCM mutations decrease the number of heads in the IHM ‘OFF’-state, which results in more heads available for interaction with actin and hypercontractility of the heart.

**Figure 1.**
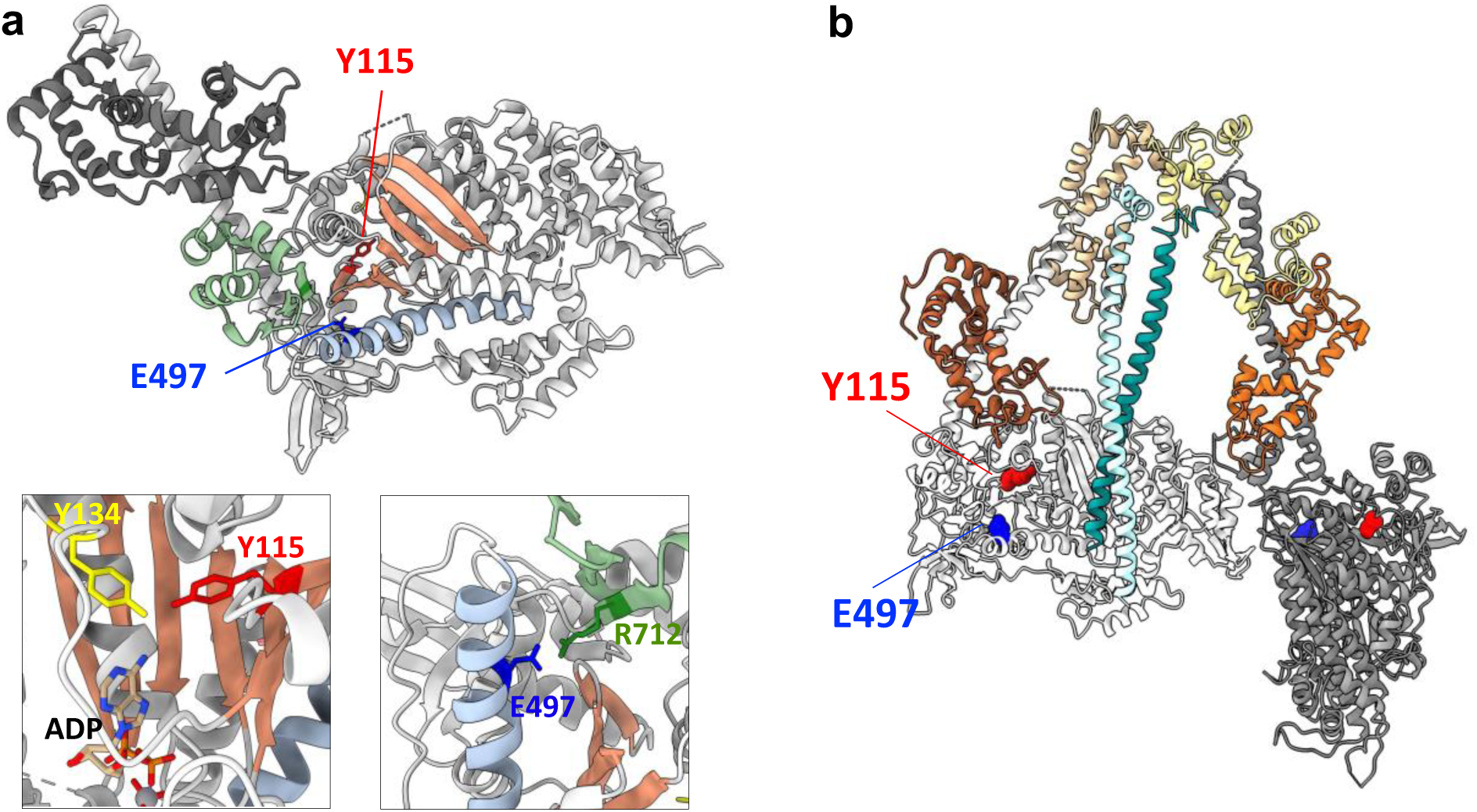
Location of the residues Y115 and E497 on myosin’s three-dimensional structure. (a) Residues Y115 and E497 are indicated on the motor head; X-ray crystal structure of bovine cardiac myosin S1 fragment in the pre-powerstroke state is used for the illustration (PDB ID: 5N69). Heavy chain (light grey), ELC (dark grey) and key relevant structural elements (central β-sheet: coral, relay helix: light blue and converter: green) are highlighted. Left panel on the bottom shows the residue Y115 (red) located in close proximity to Y134 (yellow) in the purine-binding loop and the nucleotide (ADP). Right panel on the bottom shows the interaction between E497 (blue) on the relay helix and R712 (green) on the converter. (b) Residues Y115 and E497 are indicated on the cryo-EM structure of the folded-back state of human β-cardiac myosin (PDB ID: 8ACT). Heavy chain (two shades of grey), ELC (two shades of orange), RLC (two shades of yellow) and proximal S2 tail (two shades of cyan) are indicated. Protein images were generated using *UCSF Chimera*.

## RESULTS

### Selection of HCM mutations not located at the intramolecular interfaces stabilizing the cardiac IHM

The motor function of myosin depends on allosteric communication between its nucleotide-hydrolysis site, actin-binding surface, and the lever arm, which is facilitated by several loops and deformable connectors within the motor head^44^. HCM-causing mutations linked to *MYH7* could result in hypercontractility by affecting the fundamental molecular parameters of the motor (such as intrinsic force or velocity of movement along actin) and/or by destabilizing the IHM ‘OFF’-state. For the purpose of this study, we focused on pathogenic HCM mutations that do not appear to affect residues involved in the intramolecular interactions stabilizing the cardiac IHM. After examining the high-resolution cryo-EM structure of the cardiac IHM (PDB ID: 8ACT), we chose two such mutations, Y115H and E497D (Fig. 1a, b), both of which are known to be clinically pathogenic^45,46^ but remain uncharacterized.

Y115 resides on the first strand of the conserved central seven-stranded β-sheet, lying very close to the ATP-binding pocket of myosin. Crystal structures of the β-cardiac myosin motor head (in both the pre-powerstroke state complexed with omecamtiv mecarbil, OM-PPS, PDB ID: 5N69^47^, and the post-rigor state complexed with MgADP, PDB ID: 6FSA^31^) show contacts between Y115 and the conserved purine binding loop (residues 126-134, NPXXXXXXY), specifically with residue Y134 which forms a hydrogen bond with the adenine ring (Fig. 1a). The Y115H HCM mutation might alter the nucleotide binding properties of myosin, but it is not expected to affect IHM stability, based on previous studies of the human β-cardiac myosin quasi-atomic IHM models (PDB 5TBY^32^ and MA1^31^) which are supported by the high-resolution cryo-EM structure of the cardiac IHM^39^. E497 lies in the relay helix that transmits rearrangements occurring within the motor head to the converter domain, which in turn transmits them to the lever arm that effects the power stroke. E497 forms a highly conserved salt bridge with R712 in the converter (seen in both 5N69 and 6FSA crystal structures) (Fig. 1a), and mutations at both these positions cause HCM in humans. Mechanical uncoupling of the relay-converter interface *via* disruption of this interaction in *Drosophila* indirect flight muscle myosin led to sarcomere assembly defects and impaired myosin function^48^, and more recently, the R712L HCM mutation was shown to result in a defective working stroke of human β-cardiac myosin^49^. The E497D HCM mutation is linked to a non-obstructive apical hypertrophy morphology, but this mutant variant of human β-cardiac myosin has not been extensively characterized. E497 was proposed to be involved in major IHM interactions based on the analysis of the IHM model PDB 5TBY, and a mutation at this position was predicted to impair IHM formation^32^. However, the homology model was based on a low resolution model of tarantula skeletal myosin thick filaments and the recent high-resolution structure of human β-cardiac myosin IHM reveals that E497 is in fact not part of the interactions stabilizing this conformation of cardiac myosin.

Since both Y115H and E497D mutations are expected to significantly affect the enzymatic activity of myosin by perturbing key long-range communication networks within the motor head, we first characterized the effect of these mutations on the fundamental contractility parameters of human β-cardiac myosin.

### The HCM mutations Y115H and E497D have varying effects on myosin ATPase activity and actin-sliding velocity

For this set of experiments, we used the single-headed recombinant sS1 construct of human β-cardiac myosin, consisting of the catalytic head and a part of the lever arm, bound to the essential light chain (ELC) (SDS-PAGE gels in Supplementary Fig. S1). We first examined the effect of these mutations on the actin-activated ATPase activity of myosin (Fig. 2a and Table 1). Steady-state actin-activated ATP hydrolysis rates of wild type (WT) and Y115H mutant myosin were comparable (WT, *k*_cat_ = 3 ± 0.2 s^−1^; Y115H, *k*_cat_ = 3.1 ± 0.1 s^−1^), but the mutant protein had a higher apparent actin affinity compared to WT protein (WT, *K*_app_ = 19 ± 2.4 μM; Y115H, *K*_app_ = 10 ± 0.4 μM). On the other hand, the E497D myosin had a significantly faster ATPase rate (WT, *k*_cat_ = 3 ± 0.2 s^−1^; E497D, *k*_cat_ = 4.6 ± 0.3 s^−1^) but the apparent actin affinity remained comparable to WT myosin (WT, *K*_app_ = 19 ± 2.4 μM; E497D, *K*_app_ = 17 ± 1.3 μM).

**Figure 2.**
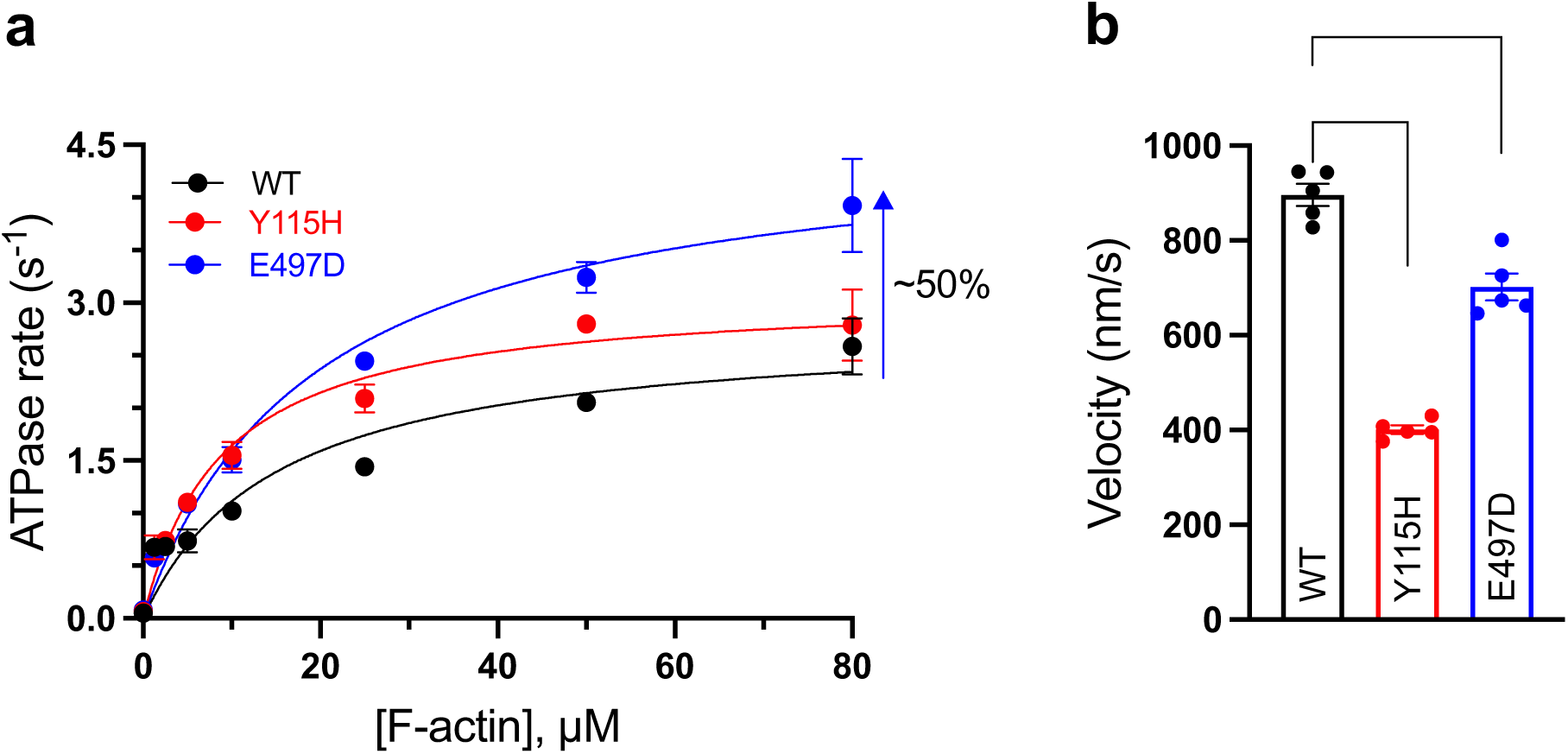
HCM mutations Y115H and E497D alter the actin-activated ATPase activity and actin sliding velocity of β-cardiac myosin sS1. (a) Steady-state actin-activated ATPase data for WT, Y115H and E497D sS1 ensembles. A representative dataset is shown from one side by side preparation of the three proteins; mean ± SD from three technical replicates at each actin concentration is plotted. The data is fit to the Michaelis-Menten equation (solid line) to determine *k*_cat_ and *K*_app_ for each myosin and the shaded area indicates the computed 95% confidence intervals. A summary of values and statistical comparisons from ATPase experiments from four independent protein preparations is provided in Table 1. (b) Actin sliding velocities for WT, Y115H and E497D sS1 myosin ensembles at 23 °C. Filtered mean velocity (MVEL_20_, see methods and Supplementary Fig. S2) measurements from 5 independent experiments across 4 protein preparations are reported, mean ± SEM is plotted; all values of velocities are given in Table 1. *** indicates p ≤ 0.001, **** indicates p ≤ 0.0001.

We next examined the effect of both mutations on myosin’s ability to move actin filaments in an unloaded motility assay. Actin-sliding velocity was significantly reduced by both mutations: ~55% by the Y115H mutation and ~20% by the E497D mutation (at 23 °C WT, *V* = 896 ± 23 nm/s; Y115H, *V* = 401 ± 9 nm/s; E497D, *V* = 702 ± 28 nm/s; Fig. 2b and Table 1, Fast Automated Spud Trekker or FAST analysis^50^ in Supplementary Fig. S2). Although Y115H myosin propelled actin filaments dramatically slower compared to WT myosin, we observed similar percentages of stuck actin filaments for the two proteins (WT: 0-5% and Y115H: 5-8%). In contrast, ~15-20% of actin filaments were immobile for E497D myosin, indicating that the mutant protein had more inactive heads compared to WT protein. Similar observations have previously been reported for the HCM mutation P710R^51^ and the dilated cardiomyopathy mutation S532P^50^. ‘Deadheading’ E497D myosin (see Methods) to eliminate molecules that irreversibly bind and stall actin filaments only modestly reduced the stuck percentage (10-12% filaments remained stuck even after deadheading), and had no effect on the measured actin-sliding velocity (Supplementary Fig. S3), indicating that the observed ~20% drop in velocity is likely a direct result of the mutation’s effect on motor function.

Overall, we observed that as expected based on their locations, both Y115H and E497D mutations significantly altered the ensemble properties of myosin. Under the detachment-limiting conditions typically employed for the motility experiments described here, decreased actin-sliding velocity by the mutations can be explained by a change in either the actin bound time and/or myosin’s step size. We next sought to determine the effect of the Y115H and E497D mutations on these parameters using single-molecule optical trapping experiments.

### Both mutations affect the load-dependent actin detachment kinetics, and, unexpectedly, the transducer mutation Y115H reduces myosin’s step size

Motor properties of single myosin molecules were determined by harmonic force spectroscopy (HFS)^52^ using the three-bead optical trap system (Supplementary Fig. S4). At physiological ATP concentrations, the rate of ADP release limits the detachment of cardiac myosin from actin, and as expected from the force-velocity relationship of cardiac muscle contraction, this rate depends exponentially on applied load. Using HFS, the lifetimes of binding events between a single myosin sS1 molecule and an actin filament were measured under a range of assistive and resistive load forces, applied randomly and automatically over the course of many binding events by harmonic oscillation of the stage upon which myosin is fixed to a bead platform for interaction with the trapped actin filament. HFS has been successfully used in the past to quantify the effects of several pathogenic mutations as well as small molecule myosin modulators on the load-dependent actin detachment kinetics of human β-cardiac myosin^37,51,53,54^.

Experimentally obtained force dependent detachment kinetics of the WT and mutant molecules were fit to the harmonic force-corrected Arrhenius equation (equation 1), as described previously^52,53,55^, to obtain detachment rates (*k_det_*), as a function of the externally applied load force (*F*):

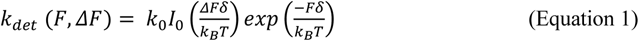

where *k*_0_ is the detachment rate at zero external force, *δ* is a measure of force sensitivity of individual molecules, *ΔF* is the amplitude of the sinusoidal force, *k_B_* is the Boltzmann’s constant, and T is the temperature. A zero-order Bessel function (*I*_0_) is used to correct for the sinusoidal nature of the applied force. *I*_0_ is a function of *ΔF*.

In agreement with the previously published values, WT sS1 myosin had a detachment rate at zero load *k_0_* = 119 ± 38 s^−1^ (Fig. 3a) and force sensitivity of the detachment rate *δ* = 1.03 ± 0.23 nm (Fig. 3b, results from 14 molecules). Y115H and E497D mutations oppositely affected the actin-detachment kinetics at zero load (Y115H *k_0_* = 63 ± 18 s^−1^, results from 12 molecules, and E497D *k_0_* = 309 ± 70 s^−1^, results from 14 molecules, Fig. 3a), while the force sensitivity was significantly different only for the E497D mutant molecules (Y115H *δ* = 1.02 ± 0.14 nm, E497D *δ* = 0.54 ± 0.15 nm, Fig. 3b). Simply put, the Y115H mutation significantly increased the average time for which myosin molecules bound to actin under zero load (~16 ms for Y115H myosin *vs.* ~8 ms for WT myosin), without affecting the load sensitivity of the bound times. On the other hand, E497D myosin molecules bound actin only for ~3 ms on average under zero load and displayed a much shallower/less sensitive dependence on applied load. Further analysis of the same HFS data was done to obtain step sizes for each interacting myosin molecule^51^, see methods. Remarkably, the active site adjacent mutation Y115H dramatically decreased myosin’s step size by ~50% (*d* = 2.2 ± 1 nm), while step sizes of WT (*d* = 4.6± 1.5 nm) and E497D (*d* = 4.7 ± 1 nm) myosin molecules were comparable (Fig. 3c). Under the experimental conditions employed here^56,57^, ensemble velocity can be approximated as the step size divided by the time myosin is strongly-bound to actin (*V = d/t_s_)*, and since *t_s_* is inversely proportional to the detachment rate, *V = d*k_det_*. We note that the changes in actin-detachment kinetics and step-sizes observed in single molecule experiments account for the change observed in ensemble motility measurements for Y115H myosin, but not for E497D myosin, which is further discussed in Supplementary Note 1.

**Figure 3.**
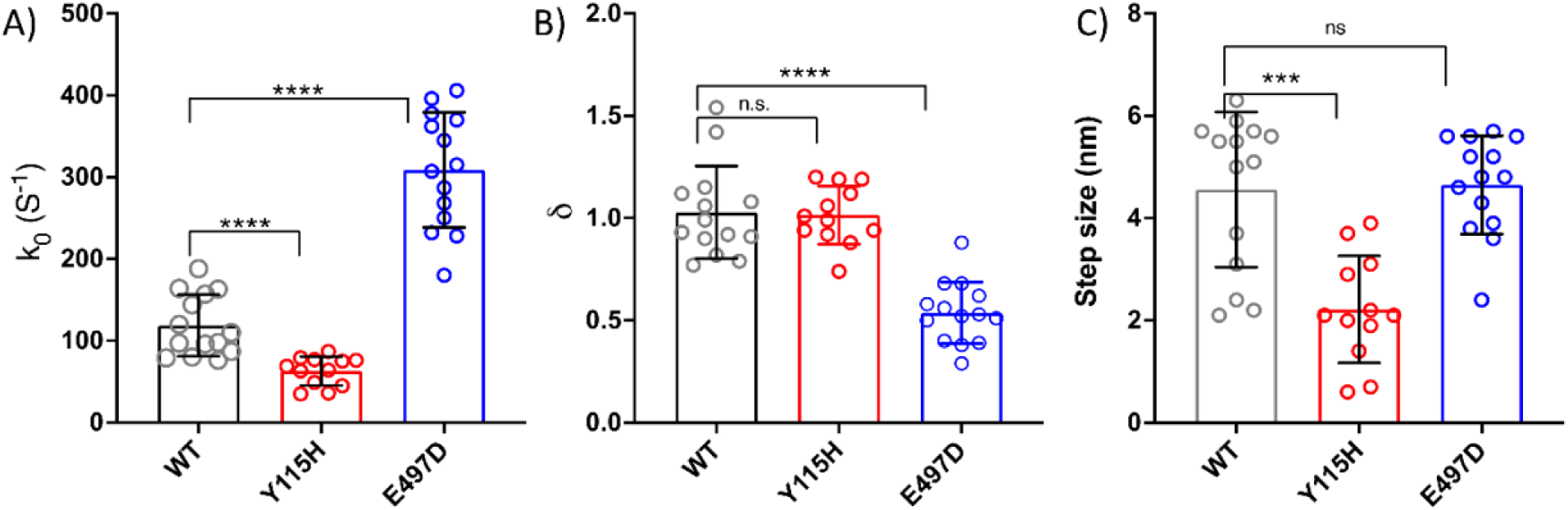
HCM mutations Y115H and E497D alter β-cardiac myosin sS1 single molecule biomechanics. The (a) actin-detachment rates at zero load (k_0_), (b) force sensitivity (δ), and (c) step-size of single molecules of WT (n = 14 molecules from 2 independent protein preparations), Y115H (n = 12 molecules from 2 independent protein preparations) and E497D (n = 14 molecules from 2 independent protein preparations) β-cardiac myosin sS1are plotted. *** indicates p ≤ 0.001, **** indicates p ≤ 0.0001.

### Single molecule and ensemble measurements do not suggest a clear mechanism of hypercontractility by the E497D mutation

The force produced by an ensemble of motor proteins should theoretically be dependent on the fraction of the time each motor spends (within its ATPase cycle) in the force-producing state. The measure of this “duty ratio r(F)” as a function of load force F, can be obtained as follows:

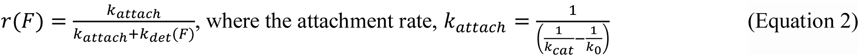

Here, *k_cat_* is the actin-activated ATPase rate measured by steady-state assay. Myosin molecules that spend less time during each catalytic cycle in the actin-bound, force producing state have lower duty ratios and faster actin detachment rates. The force-dependent duty-ratios for WT, Y115H, and E497D are plotted in figure 4a. The duty ratios for Y115H remained higher than WT for the whole force range due to lower *k_det_* (*F*) values for Y115H, as expected. The duty ratios for E497D were lower that of WT; however, near zero external force, the duty ratio for E497D remains comparable to that of WT. This difference in the force dependence of duty ratios between WT and E497D is a result of the lower force-sensitivity of the E497D mutant.

**Figure 4.**
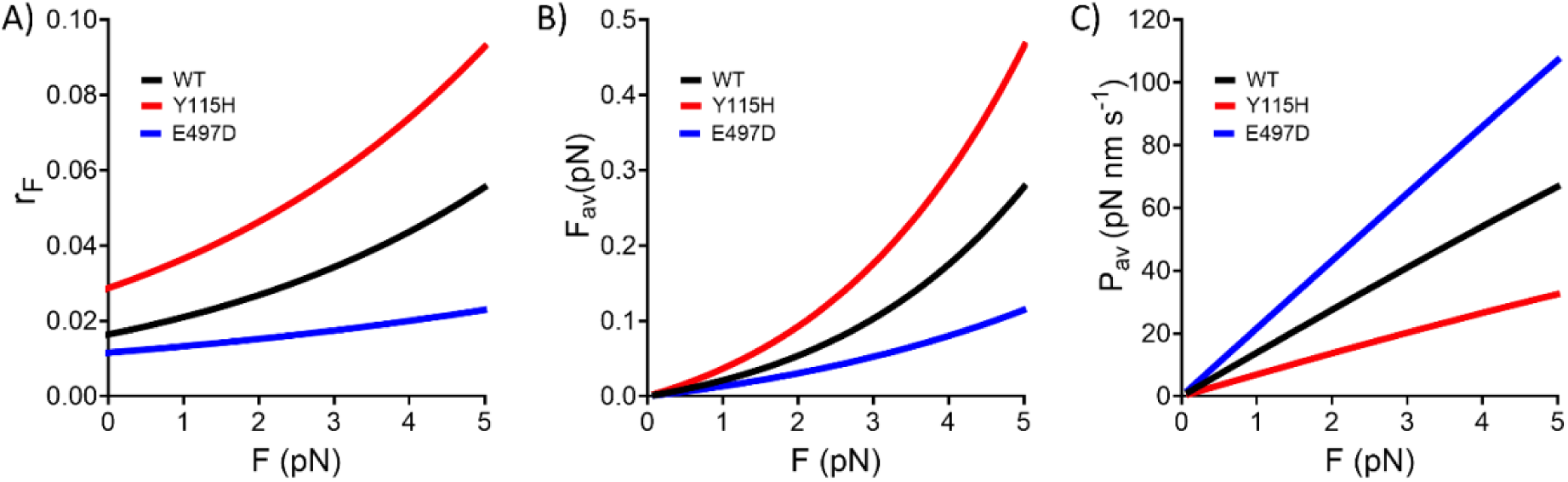
HCM mutations Y115H and E497D alter the duty-ratio, average force production, and average power production of β-cardiac myosin sS1 in opposite ways. From the measured values of detachment rate k_0_, actin-activated ATPase rate k_cat_, force-dependent detachment rate k_det_(F), and step size (d), (a) duty ratios (equation 2), (b) average force (equation 2), and (c) average power outputs (equation 4) are calculated for the resistive force-range for WT, Y115H, and E497D constructs. Curves for WT are shown in black, Y115H in red, and E497D in blue.

Using duty ratio, the average force produced by a single molecule can be calculated using the following relation.

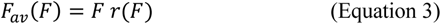

Here, *F* is the load force. The plot for the average force against different external forces follows the same trend as that of the duty-ratios (Fig. 4b). Finally, using the *F_av_*(*F*), the average power produced by a single molecule can be obtained from equation 4.

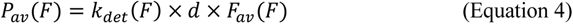

The average power produced by the myosin molecules against different load-forces is plotted in figure 4c. Overall, Y115H and E497D showed different but opposite trends compared to WT in the average power plot, with the significantly higher k_det_ of E497D leading to greater power despite its relatively decreased F_av_, while the lower k_det_ and smaller step-size of Y115H account for its lower average power.

Taken together, single molecule and ensemble experiments show that several changes occur in the biomechanical properties of the myosin motor domain in response to both the Y115H and E497D mutations. These changes result in higher predicted power output for the E497D mutation, but do not show a clear mechanism of hypercontractility by the Y115H HCM mutation. So, we next assessed the effects of these mutations on autoinhibition of myosin. For these experiments, we used purified two-headed HMM constructs containing a fragment of the proximal S2 tail that is long enough to stabilize the folded back conformation of myosin in vitro.

### Both Y115H and E497D mutations destabilize SRX myosin and increase the rate of ATP turnover by DRX myosin

An often used assay assumed to be a proxy for the IHM ‘OFF’-state is a measurement of the basal rate of ATP hydrolysis by myosin in the absence of actin using the single mant-ATP turnover (STO) assay^16^. In this pulse-chase experiment, myosin is loaded with roughly equimolar amount of fluorescent mant-ATP and then chased with excess unlabelled ATP after briefly allowing for mant-ATP hydrolysis^16,38,58^. The release of mant-ADP results in fluorescence decay that fits well to two exponential rate constants: the fast rate (~0.03 s^−1^) presumably represents the basal ATPase of myosin heads in the ‘open’ DRX state, and the slow rate (~0.003 s^−1^) arises from ‘closed’ myosin heads in the SRX state. Previous work, including ATPase assays, salt dependence of DRX:SRX ratio, EM studies, FRET studies and fiber-diffraction experiments, suggests that the biochemically measured SRX myosin in long-tailed two-headed constructs might be correlated with a folded-back inactive structural state^38,58–60^, in keeping with conventional thinking by investigators studying HCM^6,20,23,61–63^. Several HCM mutations have been shown to reduce the SRX fraction in STO assays^23,34–37,51,64^, and we applied this technique to Y115H and E497D mutant myosins.

For these measurements, we used 8-hep (short-tailed) and 15-hep (long-tailed) myosin constructs, containing the first 8 and 15 heptads respectively of the proximal S2 tail, and stabilized in the dimeric form by insertion of a GCN4 leucine zipper after the S2 fragment. Previous studies with smooth muscle myosin demonstrated that at least the first 15 heptads of the rod domain were required to form the completely inactive folded-back state of myosin^65^, and this was recently shown to be true for human β-cardiac myosin as well^39,59^. We used the previously uncharacterized 8-hep myosin construct as the uninhibited fully active control as its shorter S2 fragment should not support the formation of the folded-back state. 8-hep and 15-hep constructs are similar in principle to the previously characterized 2-hep and 25-hep constructs of myosin^38^. These new constructs were designed because 2-hep myosin has an uncharacteristically tight *K_app_* for actin, and 25-hep myosin has increased protein loss during the purification process because of a significant overlap between the ion-exchange elution profiles of 25-hep myosin and endogenous full length mouse myosin from C2C12 cells.

We first examined the turnover of mant-ATP by WT 8 and 15-hep proteins at varying salt (potassium acetate, KOAc) concentrations (Supplementary Fig. S5 and Table 2). The fluorescence decays for both the proteins were bi-exponential and the fitted fast (0.01-0.03 s^−1^) and slow rates (0.002-0.004 s^−1^) were similar to the previously measured DRX and SRX rates with WT 2 and 25-hep proteins^38^. The SRX:DRX ratio in the short-tailed WT 8-hep myosin did not display salt dependence, indicating its inability to form the folded-back state, while for the long-tailed WT 15-hep myosin the SRX fraction decreased with increasing salt concentration (Supplementary Fig. S5). These results suggest that in this case the SRX biochemical state correlates with a structural state with inhibited activity that is stabilized by ionic interactions. The IHM state is the likely structural basis for the SRX state measured here, which is also supported by our recent cryo-EM studies where the WT 15-hep protein was used (without glutaraldehyde cross-linking or stabilization by small molecules) for cardiac IHM structure determination^39^.

The relatively slow kinetics of the basal ATP turnover are typically measured by manual-mixing in a plate-based fluorimeter (15-20 seconds measurement dead-time). Unlike previously studied mutations^34–37,51^, both Y115H and E497D 15-hep proteins displayed significantly faster kinetics of mant-ADP release (Fig. 5a-c), which were captured using an automated reagent injector that dispenses and immediately starts measuring, dropping the dead-time to ~2 seconds. At low salt (5 mM KOAc), where SRX myosin is most stabilized (Supplementary Fig. S5a), we found that both Y115H and E497D mutations dramatically reduced the percentage of molecules in the SRX state (Fig. 5d and Table 2). ~55% of WT 15-hep myosin hydrolysed ATP at the slow rate, but only <10% molecules in both Y115H 15-hep myosin and E497D 15-hep myosin were estimated to be in the SRX state. The percentage of molecules in the SRX state was not significantly different between the WT, Y115H and E497D 8-hep proteins (Fig. 5d). Both Y115H and E497D mutations significantly increased the fast rate compared to WT protein (Fig. 5e), in both 8-hep and 15-hep backgrounds (15-hep *k_fast_*: WT = 0.015 ± 0.001 s^−1^, Y115H = 0.05 ± 0.003 s^−1^, E497D = 0.07 ± 0.003 s^−1^, see Table 3 for 8-hep rates). The SRX rate was not affected by these mutations (Fig. 5e).

**Figure 5.**
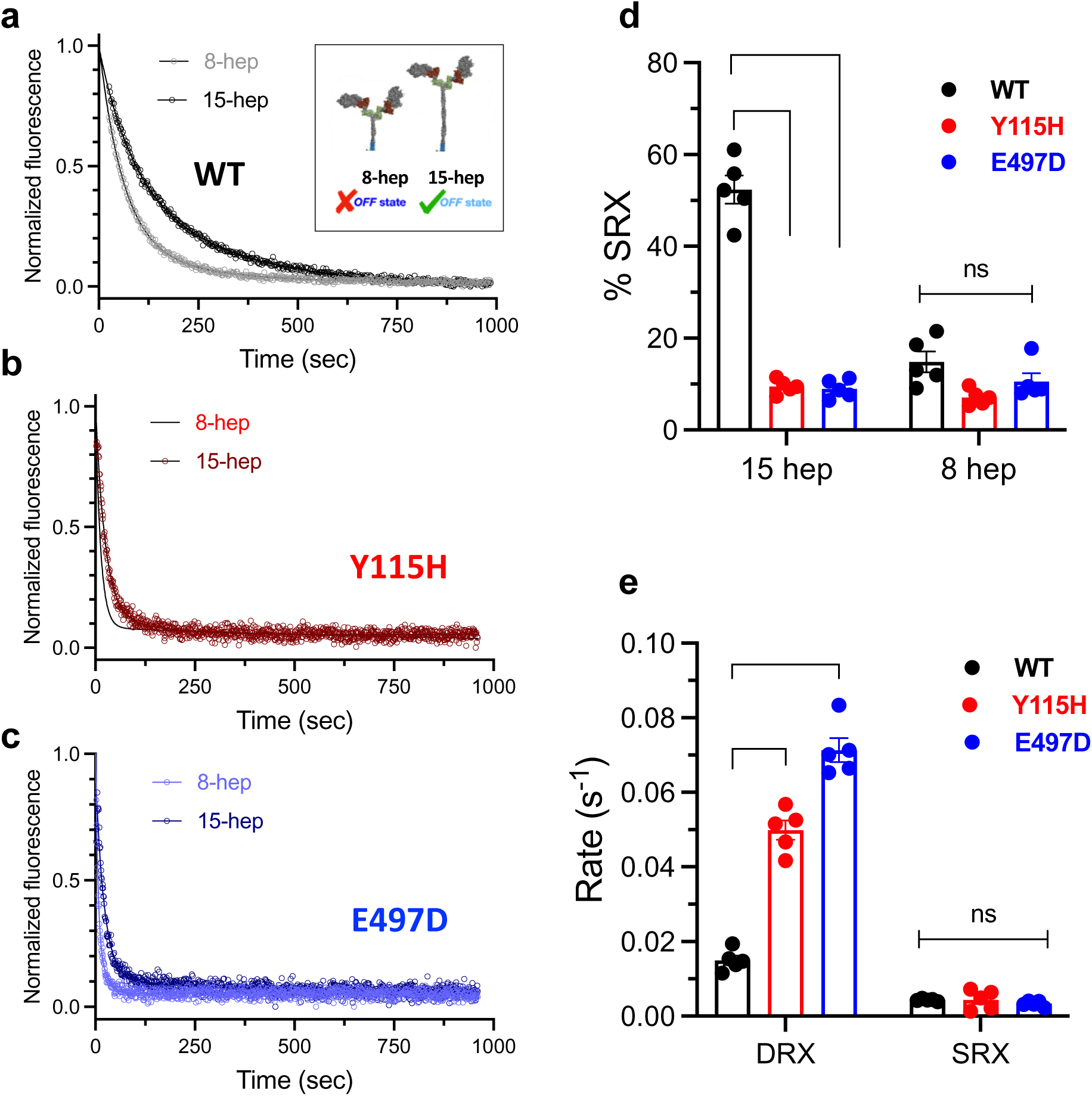
HCM mutations Y115H and E497D reduce the super relaxed state fraction of β-cardiac myosin HMM. (a-c): Representative traces showing the kinetics of mant-ATP turnover by 8 and 15-hep constructs of (a) WT, (b) Y115H, and (c) E497D myosin. The inset in (a) shows a schematic of two-headed short (8-hep) and long (15-hep) tailed HMM protein constructs. The data is fit to a double exponential (solid line) and fitted fluorescence values at t = 0 and t = ∞ were used to normalize the individual curves. (d) Quantification of the relative amplitude of the slow rate constants (%SRX). Data is combined from multiple florescence transients from 3 independent protein preparations, mean ± SEM is plotted. (e) Fast (DRX) and slow (SRX) rate constants for 15-hep constructs (see SI for 8-hep rate constants for all three proteins). (**** indicates p ≤ 0.0001) All values are given in Table 2.

### The HCM mutations Y115H and E497D increase the number of myosin heads available for actin binding

While SRX measurements have been used as a reasonable proxy for the IHM ‘OFF’-state, they are indirect and even single-headed S1 preparations show a fraction of SRX heads in a STO experiment^38^, showing that SRX heads are not always correlated with the IHM ‘OFF’-state heads^25,66–68^. Furthermore, the lever arm mutations D778V, L781P and S782N do not show a change in SRX population, while direct measurements of an increase in active heads using actin-activated ATPase rate (*k_cat_*) measurements using the Long-tail/Short-tail ATPase Ratio Assay (LSAR) clearly show an increase in the number of active heads caused by those lever arm mutations^37^.

To confirm that the Y115H and E497D mutations increase the number of active heads in a two-headed myosin population capable of forming the IHM ‘OFF’-state, we used the LSAR assay. Similar to previous observations with the WT 25-hep (long-tailed) and 2-hep (short-tailed) myosin constructs^38^, we measured a significant ~40% drop in the *k_cat_* of WT 15-hep (long-tailed) myosin compared to WT 8-hep (short-tailed) myosin (WT 8-hep, *k*_cat_ = 4.1 ± 0.2 s^−1^; WT 15-hep, *k*_cat_ = 2.6 ± 0.1 s^−1^; *P* = 0.0016; Fig. 6a and Table 2). These results confirm that, as anticipated, a large fraction of the myosin population in WT 15-hep myosin is in the folded-back state, which decreases the number of myosin heads that bind to actin and quickly release ATP hydrolysis products.

**Figure 6.**
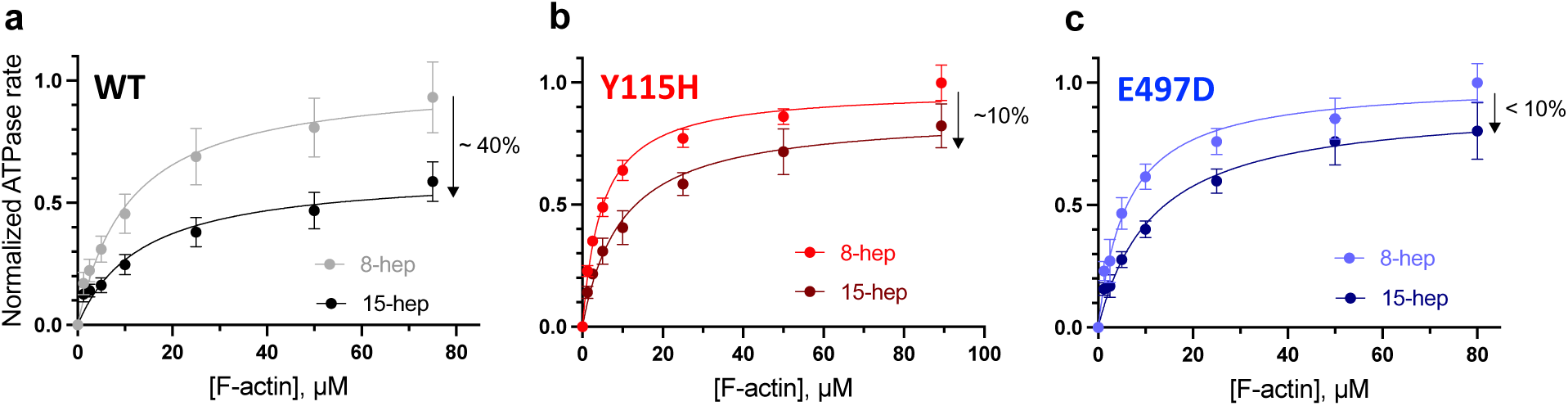
HCM mutations Y115H and E497D increase the number of active β-cardiac myosin heads as measured by the actin-activated ATPase LSAR assay. Actin-activated ATPase curves for 8-hep and 15-hep constructs of (a) WT, (b) Y115H, and (c) E497D myosin. Raw data is combined from 12 experimental replicates from 4 independent protein preparations for WT and 9 experimental replicates from 3 independent protein preparations for mutants. Each data point represents the average across all replicates (mean ± SEM is plotted). ATPase curves in each panel are normalized to the respective 8-hep *k*_cat_ (see Table 2 for raw values and statistics). Solid lines are the fitted lines (Michaelis-Menten kinetics) and shaded areas indicate the 95% CI of the fit. Downward arrows indicate the average % drop in *k*_cat_ of 15-hep constructs relative to 8-hep controls.

Previous experiments have shown that a number of HCM mutations significantly reduce the ~40% drop in the *k_cat_* of WT long-tailed myosin (25-hep) compared to WT short-tailed myosin (2-hep), in keeping with a decrease in the population of IHM ‘OFF’-state myosin molecules and a resulting increase in the number of heads available for interaction with actin^34–37,51^. This LSAR assay is the most direct method for measuring an increase in the number of heads available for interaction with actin. We therefore measured the ATPase activity of Y115H and E497D myosins using the 8-hep and 15-hep constructs. For each mutant, a set of WT 8-hep, mutant 8-hep and mutant 15-hep proteins was grown and processed for purification in parallel to reliably capture the differences in *k_cat_* due to the mutations. Firstly, consistent with the trends observed with sS1 proteins (Fig. 2a and Table 1), ATPase assays with 8-hep myosin showed WT and Y115H mutant as having similar activity while the E497D mutation showed a ~50% increase in *k_cat_* (Table 2). Remarkably, we found both the Y115H and E497D mutations reduced the ~40% drop in ATPase seen between the 15-hep and 8-hep WT protein to <10% (Fig. 6b, c). There was no significant difference between the mutant 8-hep and mutant 15-hep myosin ATPase rates for either mutant protein: for Y115H, 8-hep *k_cat_*: 4.3 ± 0.2 s^−1^ and 15-hep *k_cat_*: 3.8 ± 0.3 s^−1^ (*P* = 0.16), and for E497D, 8-hep *k_cat_*: 6.7 ± 0.1 s^−1^ and 15-hep *k_cat_*: 6.1 ± 0.3 s^−1^ (*P* = 0.34) (mean ± s.e.m. calculated from data from three independent protein preps with 3 replicates each is reported for each mutant). This suggests that both the Y115H and E497D mutations significantly impair myosin’s ability to form the IHM ‘OFF’-state, leading to increased availability of myosin heads for actin binding and force production. We note that the basal ATPase rates (extracted from the zero actin well) for the Y115H and E497D proteins (both 8-hep and 15-hep constructs) were elevated compared to WT protein (Supplementary Fig. S6), correlating with the increased DRX rates for the mutants. For all 6 proteins (WT, Y115H and E497D 8-hep and 15-hep constructs), the measured basal rates corelated well with the amplitude weighted sum of the DRX and SRX rate constants for each protein (calculated basal rates, Supplementary Fig. S6).

### Molecular dynamics simulations

We carried out MD simulations to understand the long-range perturbations in response to Y115H and E497D mutations that result in impaired autoinhibition of myosin. We chose to simulate the pre-powerstroke conformation (Fig. 7a) of myosin. In the pre-powerstroke conformation, myosin S1 heads are primed to interact with thin filaments and are characterized by a rocking motion of the lever arm (monitored here by residues 767-810) and ELC relative to the N-terminal domain. The interface between the converter domain and relay helix serves as a fulcrum point for this rocking motion. In the WT simulations, the tail and N-terminal domain sample diverse conformational states that interconvert on timescales ≤ 500 ns (Fig. 7b). We characterized this motion by clustering simulations on the C_α_ atoms of residues 96-110 (an N-terminal domain α-helix), 155-166 (an N-terminal domain α-helix), 473-503 (the relay α-helix), and 769-783 (the tail α-helix) (Fig. 7c). The structures depicted in Figure 7c represent the most frequently observed motor domain:tail orientations. While the range of orientations accessed by the WT and mutant myosins was similar, the frequency of sampling was not. We also characterized this motion by calculating specific interhelical angles between the tail and either the relay helix (Fig. 7d), the 96-110 α-helix (Fig. 7e), or the 155-166 α-helix (Fig. 7f). Relative to the WT simulations, the E497D simulations and – to a lesser extent the Y115H simulations – modified the dynamics of the motor domain/tail interface. The mutations did not appreciably change the conformations explored by the converter domain and tail; instead they modified the relative frequency at which different conformations were sampled (Fig. 7c). The distributions of interhelical angles suggest that the E497D mutation increased conformational heterogeneity among the motor domain, converter and tail whereas the Y115H mutation decreased it (Fig. 7d).

**Figure 7.**
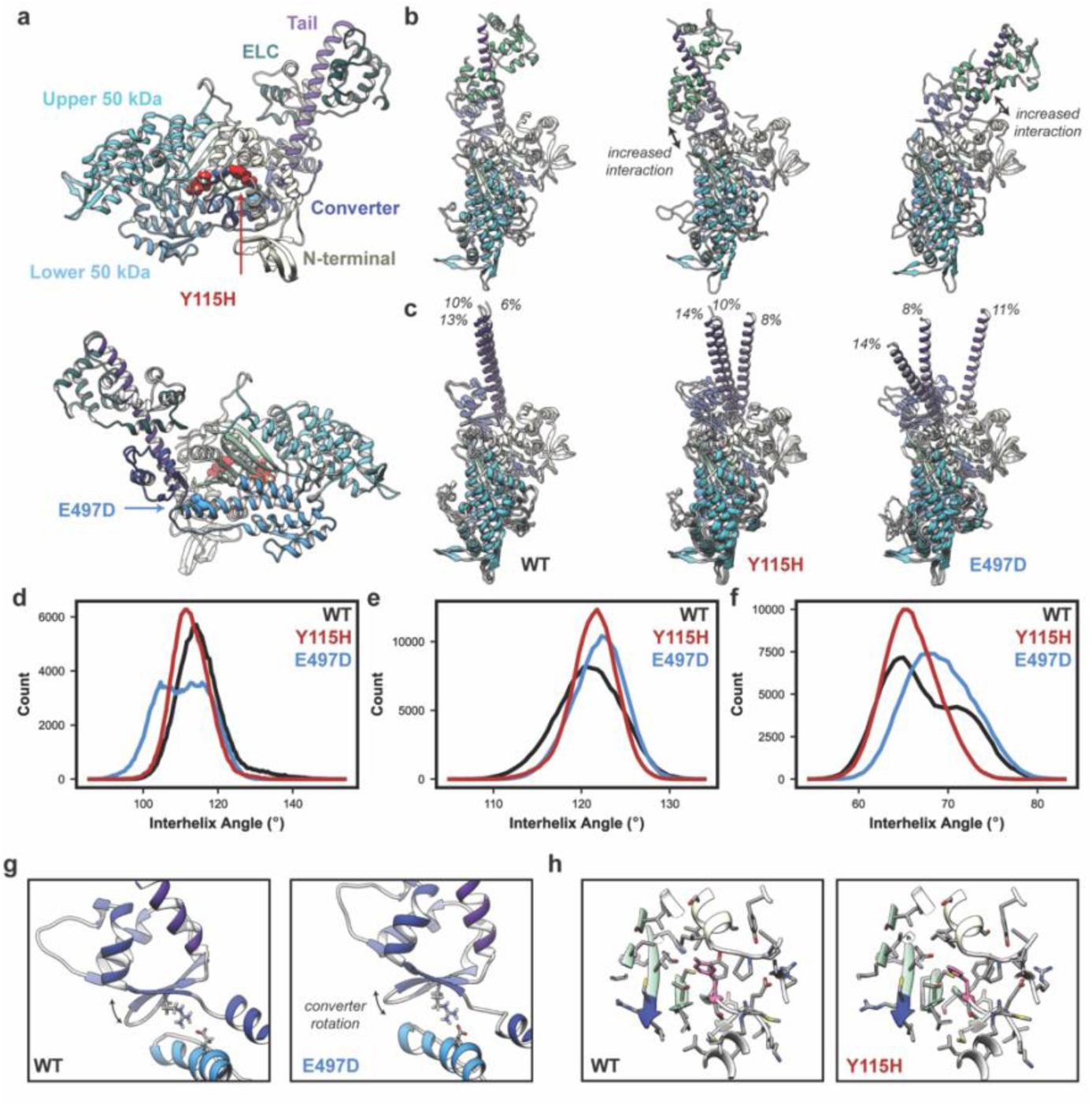
Y115H and E497D modified motor-domain : lever arm dynamics in molecular simulations of pre-powerstroke myosin. (a) Our molecular model of the pre-powerstroke (ADP.P_i_) state of β-myosin. WT Y115 and E497 are shown as red and blue spheres, respectively. (b) The converter domain and tail explored multiple conformers relative to the motor domain. The X-ray like (and most commonly observed) conformer is shown on the left. Less frequently observed conformers were associated with increased interactions between the tail/ELC and N-terminal domain. (c) The top 3 most highly populated clusters are shown and the frequency at which each cluster was observed in the ensemble is annotated. Interhelical angles between the tail and (d) the relay helix, (e) 96-110 α-helix, or (f) the 155-166 α-helix. Snapshots taken at the end of the MD simulations highlight the converter domain rotation observed in the E497D simulations (g) and the relative stability of the Y115H mutation site (h).

Next, we analysed local structural dynamics of the WT and mutation simulations to understand the molecular origins of the observed behaviour. In the WT simulations, E497 formed a salt bridge with R712 (Fig. 7g, left panel). This salt bridge was maintained in E497D simulations (Fig. 7g, right panel). However, the shorter length of the Asp side chain required a rotation of the converter domain to maintain the salt bridge, which may ultimately modify the relative orientation of the tail and motor. In the WT simulations, Y115 is buried in the central β-sheet and forms numerous interactions with other residues in the core (Fig. 7h). In the Y115H simulations there were subtle changes among side-chain to side chain interactions in this region, though the overall conformation remained similar to the WT, and H115 retained a WT-like rotameric state (Fig. 7h). Though this region is 2.3 nm away from the motor domain/converter domain interface (near the active site), the small magnitude changes in structure that were observed did not suggest obvious molecular mechanisms that relate to changes in the tail or converter domain.

## DISCUSSION

Experimental support for the central role played by myosin autoinhibition in the regulation of cardiac contractility, and the association of its dysregulation with cardiomyopathies has grown tremendously in the past decade^6,22,69^. Many HCM-causing mutations appear to affect the interfaces stabilizing the cardiac IHM structure^8,11^. Aligned with these observations, a direct measure of increase in the number of functional myosin heads available for interaction with actin using the LSAR assay and/or a decrease in the SRX fraction has been observed for >15 of these when tested in vitro^34–37,51^. Not surprisingly, these correlations have led to more detailed analyses of HCM variant locations using homology modelled structures of cardiac IHM^24,31,32^, which have predicted that the pathophysiology of ~30-50% of the known pathogenic HCM mutations may be linked to their direct effects on the stability of the autoinhibited state of myosin. Here we show that Y115H and E497D mutations, which are not expected to directly affect the IHM structure, completely destabilize the SRX myosin and increase the *k_cat_* of long-tailed myosin constructs. These are not isolated findings - G256E^34^ and P710R^51^ HCM mutations also appear to interfere with myosin autoinhibition without directly altering the IHM interfaces. Altogether, these observations show that due to the extremely allosteric nature of myosin, limiting our analyses to interfacial residues on a static structure might in fact have resulted in an underappreciation of how widespread this mechanism of disease manifestation might be. Importantly, these results highlight that it is not trivial to predict the effects of a pathogenic mutation on myosin’s enzymatic activity or conformational dynamics. Purified mutant forms of bonafide human β-cardiac myosin must be analysed, preferably by the direct LSAR assay.

Unlike several previously studied HCM mutations^70^, Y115H and E497D mutations significantly affected several parameters measuring motor function. E497D increased *k*_cat_ by ~50%, which is one of the largest changes measured for human β-cardiac myosin disease variants and reminiscent of the early-onset HCM mutations D239N (50%) and H251N (24%)^71^. But the combination of the observed changes for both Y115H (unchanged *k*_cat_, reduced *k*_det_ and step-size) and E497D (elevated *k*_cat_ and *k*_det_, unchanged step-size) had opposite effects on the calculated average force and average power produced by a single myosin molecule. Overall, these changes together fail to predict a clear mechanism of hypercontractility for both Y115H and E497D. We highlight two unexpected findings from these studies. First, based on the position of Y115H, a reduction in step-size is an unexpected change. Previously, different mechanisms have been proposed to explain step-size changes observed for other mutations based on their location: uncoupling of the biochemical and mechanical activities of myosin (P710R^51^ and R712L^49^ in the converter), destabilization of the lever arm (L781P^37^), and altered lever arm priming in the pre-powerstroke state (R671C^54^ in the transducer). Judging from the location of Y115H in the motor head, it seems unlikely that this mutation affects lever arm coupling or compliance. Second, we note that E497D and R712L HCM mutations, both affecting the relay-converter E497-R712 ionic interaction, have dramatically different effects on motor function. Disruption of the E497-R712 salt bridge by charge reversing (E496R in *Drosophila* indirect flight muscle myosin^48^) or charge neutralizing (R712L in human β-cardiac myosin^49^) mutations reduces motility velocity (~70-80% drop) and myosin’s working stroke amplitude (4-fold reduced for R712L). The E497D mutation retains the negative charge but replaces Glu by the less bulky Asp residue, and it is unclear if the shorter side chain of Asp disrupts the ionic interaction with R712 or induces an abnormal conformation of the motor to allow for the formation of this interaction. The relatively subtle change in velocity (~20% drop) and lack of change in step-size for E497D as compared to mutations that clearly disrupt this salt bridge supports the latter view. Our simulations support this view (Fig. 7g) as well, but a high resolution structure of the E497D myosin (and Y115H) should be determined to highlight how these mutations can affect the structure, flexibility and stability of the actin and ADP bound state. Incidentally, a previous study^49^ had found that the E497D mutant variant of human β-cardiac myosin propelled actin filaments nearly as fast as WT protein (1.41 ± 0.19 μm/s for E497D myosin *vs.* 1.46 ± 0.11 μm/s for WT myosin). Previous experiments were done at a higher temperature (32 °C as opposed to 23 °C in the current study) using HMM constructs (sS1 constructs used here), which might result in more myosin heads on the surface, potentially contributing to the observed discrepancy.

The single consistent hypercontractile change across Y115H and E497D mutations appears to be the increased availability of active myosin heads for interaction with actin (Fig. 5 and 6). The cryo-EM structure of human β-cardiac IHM shows that both myosin heads adopt the classical pre-powerstroke state (PPS) - their lever arms are primed and their active sites are closed^39^. A pathogenic mutation can therefore destabilize the IHM either structurally (directly by altering a residue at the interface, or indirectly by being close enough to induce changes in the dynamics or flexibility of the loops and connectors and other structural elements of the motor that are engaged at these interfaces) or by affecting the stability of the PPS by altering long-range interactions that maintain the lever arm in the primed position^31^. A previous analysis of 178 pathogenic HCM mutations using the MA1 IHM model predicted ~65% of them to disrupt the IHM state, and roughly half of them were proposed to do so by destabilizing the PPS conformation^31^. In the current study, the fast rate of mant ADP release - the DRX rate in single turnover assays-was found to be significantly faster than the WT protein for both Y115H and E497D myosin (Fig. 5). During the motor cycle of myosin, ATP hydrolysis by the catalytic head is rapid, after which it is in the PPS conformation which has a weak affinity for actin, a primed lever arm, and tightly coordinated MgADP and inorganic phosphate (P_i_)^44^. In the absence of actin-binding induced conformational changes in myosin under basal turnover conditions, the release of the ATP hydrolysis products is quite slow, and P_i_ release limits the release of MgADP. Since the stability of the PPS state is linked to the trapping of Pi and MgADP, the 3-5 fold increase observed in the fast rate of mant ADP release for the mutants suggests that both Y115H and E497D mutations likely destabilize the PPS conformation of myosin, which in turn diminishes their ability to adopt the folded-back state. The ATP binding rates for both mutants were found to be very similar to WT myosin (Supplementary Fig. S7), and these data suggest that it is the destabilization of the pre-powerstroke state that leads to speeding up of the motor basal activity. Together, these suggest that the Y115H and E497D HCM mutations increase the availability of functional myosin heads for actin interaction by destabilizing the PPS conformation, which interferes with the formation of the folded-back state. Interestingly, the G256E and P710R mutations have also been proposed to destabilize the PPS conformation^31^, which is likely the reason for the decrease in autoinhibition measured experimentally for both^34,51^. In fact, an elevated DRX rate has also been measured for the P710R (modest elevation) and P710H (~3-fold, unpublished results) mutations.

We utilized molecular dynamics simulations to probe atomic mechanisms by which Y115H and E497D exert their effects. Our simulations suggest that both mutations may modify the distribution of angles formed between the tail and the motor domain in the primed state: the E497D distributions were broader than the WT and the Y115H distributions were narrower than WT. The Y115H simulations suggested that the mutation may dampen conformational motions in the tail – the interhelical angle distributions were narrower than in the WT simulations. However, there were not obvious structural changes that explained this behaviour. Rotations and structural changes to buried aromatic residues are known to occur on microsecond to millisecond timescales^72^ and simulations longer than 500 ns (as performed here) may be necessary to capture the effects of the Y115H mutation. Our simulations show evidence of conformational heterogeneity within the tail; however, the distribution of tail conformations did not converge, precluding a statistically meaningful analysis. These simulations predict that the Y115H and E497D mutations may distinctly impact the orientation of the lever arm in the pre-powerstroke state. The mutations may similarly disrupt lever arm:motor domain conformations in other chemomechanical states. The conformations accessible to the tail are complex and impact many structural and kinetic features of myosin including the stability of the interacting heads motif and actomyosin powerstroke. We note that the simulations described here are subject to several limitations, and that structural studies will be necessary to precisely describe the changes in the PPS of these mutants and how this can lead to premature product release. First, we have only explored one of many conformational states of myosin – and the extent to which mutation effects in one state extend to the others is uncertain.

Second, the simulations performed here capture changes that occur on sub-microsecond timescales. Much higher sampling is required to determine the equilibrium tail distributions and the contributions of different residues to the stability of motor domain:lever arm conformations. Finally, our simulations were initiated from homology models using a WT bovine structure bound to omecamtiv mecarbil (a small molecule that influences motor domain:tail orientation^47^) as the template and therefore must transition from a bovine WT-like ensemble to the human WT or mutant ensembles and our analyzed results likely include data from this transition period. Despite these limitations, our simulations predict that the mutations may impact the orientation of the motor domain and tail. Based on where Y115 and E497 are localized, and the data presented here, we predict that structural studies in the future may find that, (1) lack of stability of the hydrolysis products in the active site for Y115H leads to premature P_i_ release in the absence of actin due to more flexibility/destabilization of the PPS and, (2) in a distal site, the E497D mutation likely leads to more flexibility of the converter/lever arm orientation and of the relay, which also leads to a lack of stability of the products in the active site.

Finally, HCM is clinically characterized by increased energy consumption^73^, and the energy deficits recognized in HCM patients become evident before the morphological changes associated with the disease become apparent^7^. This deficit appears to be a result of altered cellular metabolism, reduced ATP synthesis due to mitochondrial dysfunction and increased ATP utilization due to a shift in the DRX⇔SRX equilibrium^7^. HCM mutations that reduce SRX proportions lead to increased ATP consumption. Interestingly, the Y115H and E497D not only disrupted the SRX rate, but also increased the DRX rate by 3 and 5-fold, respectively. These measurements suggest that the HCM mutations can have more profound direct effects on ATP consumption in cases where, in addition to disrupting the energy conserving SRX state that increases the number of heads in the DRX state, they also cause an increase in the ATP turnover rate of these DRX heads.

In conclusion, the results we present here highlight that HCM mutations away from the IHM interfaces can allosterically disrupt myosin autoinhibition. The effects of such mutations on the long-range interactions are expected to be position- and mutation-specific and can be discerned unambiguously only from a high-resolution structure of the mutant. More importantly, our results highlight that, unlike HCM mutations that directly lie at the IHM interfaces, predicting the indirect or allosteric effect of an HCM mutation on the stability of the IHM is not straightforward. The effects of HCM mutations on myosin’s regulation by autoinhibition are nuanced and may be evident only upon experimental characterization, as for the Y115H mutation (previously predicted to affect motor activity but not IHM stability^31^). A previous study suggested that IHM structural models be leveraged to improve molecular prediction of clinically actionable human variants because they found that myosin variants with unknown significance (VUS) that altered residues participating in IHM interfaces (IHM+) were associated with higher rates of adverse clinical outcomes compared to variants that did not affect IHM stability (IHM-)^23^. However, our results show that an accurate classification of a variant as IHM-is not entirely possible due to the indirect and remote effects of mutations on IHM stability that are not easily predicted.

## MATERIALS AND METHODS

### Protein expression and purification

Three different recombinant human β-cardiac myosin constructs were used for WT and mutant proteins in this study: single-headed sS1 (res 1-808), two-headed short-tailed 8-hep HMM (res 1-893) and two-headed long-tailed 15-hep HMM (res 1-942). Myosin heavy chain (*MYH7*) was co-expressed with the human ventricular essential light chain (*MYL3*) carrying an N-terminal FLAG tag cleavable by TEV protease using adenoviruses in murine C2C12 myoblasts (ATCC) differentiated into myotubes; adenoviruses were generated in HEK293T cells (ATCC) using the AdEasy Vector System (Qbiogene Inc, Carlsbad, CA, USA) and purified by cesium chloride gradient centrifugation. *MYH7* constructs contained a C-terminal GSG-RGSIDTWV affinity tag, and a GCN4 leucine zipper placed immediately before the C-t tag ensured dimerization in the two-headed constructs. C2C12 cells were grown in 15 cm dishes (~10 dishes per construct) and infected with *MYH7* and *MYL3* adenoviruses 48 hours post initiation of differentiation into myotubes. Cells were harvested into harvesting buffer (20 mM Imidazole, pH 7.5, 50 mM NaCl, 20 mM MgCl_2_, 1 mM EDTA, 1 mM EGTA, 10% sucrose, 3 mM ATP, 1 mM DTT, 1 mM PMSF and Roche Protease Inhibitors), 1 ml per dish, 96 hours post infection using a cell scraper, and immediately flash frozen in liquid nitrogen (LN_2_).

All steps for protein purification were carried out at 4 °C (on ice or in a cold room). Cell pellets were thawed and supplemented with 3 mM ATP, 1 mM DTT, 1 mM PMSF, Roche Protease Inhibitors (1 tablet per 50 mL) and 0.5% Tween-20. Cells were lysed by 50 strokes of a dounce homogenizer and the lysate was incubated for 20-30 mins to allow for the assembly of contaminating endogenous full length (FL) mouse myosin from C2C12 cells into filaments (aided by the low NaCl and high MgCl_2_ content of the harvesting buffer), which pelleted down in the subsequent spin at 30,000 rpm for 30 mins at 4 °C. The clarified supernatant was incubated with anti-FLAG resin for 1-2 hours to capture β-cardiac myosin protein, followed by washing away non-specifically bound proteins by centrifugation using wash buffer (20 mM Imidazole pH 7.5 containing 150 mM NaCl, 5 mM MgCl2, 1 mM EDTA, 1 mM EGTA, 1 mM DTT, 1 mM PMSF, 3 mM ATP, 10% sucrose and Roche protease inhibitors). For sS1 constructs, myosin was then dissociated from the resin by overnight incubation with TEV protease, cleaving the FLAG tag on ELC. For 8 and 15-hep constructs, protein-bound resin was further incubated with regulatory light chain (RLC) depletion buffer (20 mM Tris-HCl, pH 7.5, 200 mM KCl, 5 mM CDTA, pH 8, 0.5% Triton-X-100 and 1 mM ATP) for 75 mins to dissociate endogenous mouse RLC. Resin was extensively washed with wash buffer before incubating it with 10-fold molar excess His-tagged human RLC (purified from *E. coli* as described previously^24^) for 2-3 hours. Excess unbound RLC was washed away and the resin was incubated with TEV protease overnight, cleaving the FLAG tag on ELC and His tag on RLC. The next morning, dissociated myosin (sS1 or 8-hep or 15-hep) in the supernatant was collected by spinning down the resin and the protein was further purified (to remove TEV protease and endogenous mouse FL myosin) by anion exchange chromatography using a 1 mL HiTrap Q HP column (Cytiva) attached to an ÄKTA FPLC system. Myosin typically elutes at ~250 mM NaCl; fractions were pooled based on SDS-PAGE analysis and concentrated using Millipore Amicon® ultra centrifugal filters (50 kDa MWCO for sS1, 100 kDa MWCO for 8 and 15 hep) to a concentration of 10-20 μM which was determined using the BIO-RAD Quick Start Bradford assay kit. Freshly prepped protein was directly used for ATPase assays, and aliquots were flash frozen in LN_2_ for later use for in vitro motility and optical trapping experiments.

### Actin-activated ATPase

Steady-state actin-activated ATP turnover rates were measured using a plate based NADH-coupled assay as previously described^37,74^. G-actin was prepared^75^ from acetone powder from bovine cardiac LV tissue and polymerized to F-actin by dialyzing 5x into assay buffer containing 10 mM imidazole, pH 7.5, 5 mM KCl, 4 mM MgCl_2_ and 1 mM DTT. F-actin aliquots (130-160 μM) were flash frozen in LN_2_ for later use. F-actin, thawed on ice, was thoroughly mixed with 1:100 molar ratio of gelsolin (purified as described previously from *E. coli*^76^) and incubated on ice for ~30 mins. In a 96-well clear plate and 100 μL reaction volume, the actin-gelsolin mixture was diluted in assay buffer to achieve 8-10 dilutions in the 0-80 μM concentration range, and mixed with freshly prepped myosin to achieve 25 nM final concentration (250 nM in the zero actin well to measure basal ATPase rate). Actin and myosin were thoroughly mixed by constant shaking for 10 mins in a microplate reader at 23°C, following which the ATPase reaction was initiated by carefully pipetting in 20 μL of a 5X coupling solution containing lactate dehydrogenase (100 U/mL), pyruvate kinase (500 U/mL), phospho(enol) pyruvate (2.5 mM), ATP (10 mM) and NADH (2 mM). Any air bubbles were quickly eliminated by tapping with a needle and the plate was shaken for additional 5 mins before reading absorbance at 340 nm every 15-30 sec for 25 mins at 23°C. The concentration of ADP produced was calculated from the absorbance values using an ADP standard curve (0-300 μM) set up on the same plate. ADP produced over time in each reaction well was plotted and the slope of the plot (rate) was divided by myosin concentration in that well and plotted against actin concentration (after subtracting the basal rate from zero actin well from the rate at each actin concentration). ATP turnover rates (*k*_cat_) and the apparent affinity for actin (*K*_app_) were determined by fitting the data to the Michaelis Menten model of enzyme kinetics. For sS1 constructs, 3 technical replicates of each actin concentration were set up on a single plate for WT and two mutants. For HMM constructs, WT 8-hep control was always assayed together with mutant 8 and 15-hep proteins in 3 technical replicates on a single plate. Data from 3 or more independent protein preparations with 3 technical replicates in each assay were averaged (mean ± SEM are reported here). C2C12 cells expressing proteins assayed on a single plate were grown and processed in parallel to improve the robustness of these measurements.

### In vitro motility

Motility assay was carried out as described previously^50^. 5-7 flow channels were created on a glass slide using double-sided tape and #1.5 coverslips coated with 0.1% nitrocellulose/0.1% colloidin in amyl acetate. Myosin molecules were tethered *via* their C-terminal affinity peptide tag onto nitrocellulose coated coverslips before flowing in fluorescently labelled F-actin filaments. Briefly, 5-6 μL of the following solutions were sequentially flown into the channels and incubated as noted: SNAP-PDZ for 2 mins (3 μM in assay buffer containing 25 mM Imidazole pH 7.5, 25 mM KCl, 4 mM MgCl_2_, 1 mM EGTA and 10 mM DTT; SNAP-PDZ was purified from *E. coli* as described previously^50^), blocking buffer for 2 mins (assay buffer with 1 mg/ml BSA, ABBSA), myosin twice for 2 mins (~200 nM in ABBSA), blocking buffer for 1 min and final GO buffer (2.5 nM TMR-phalloidin labelled F-actin, 2 mM ATP and an oxygen-scavenging system containing 0.4% glucose, 0.11 mg/ml glucose oxidase and 0.018 mg/ml catalase in ABBSA). Slides were typically imaged within 20 mins of flowing in the GO buffer on a Nikon Ti-E inverted microscope with a 100x TIRF objective on an Andor iXon + EMCCD camera. Three movies were acquired per channel; images were taken at 2 Hz for 30 sec with 300 ms exposure using the 561 nm laser channel. Velocity measurements are extremely temperature sensitive – to minimize the error in measurements arising from temperature fluctuations, objective temperature was maintained at 23 ± 0.3 °C as closely as possible. Filament tracking and velocity analysis was performed by analysing the movies using FAST (Fast Automated Spud Trekker^50^) using the following parameters: window size *n* = 5, path length *p* = 10, percent tolerance *pt* = 20, and minimum velocity for stuck classification *minv* = 80 nm/s. Filtered mean velocity (MVEL_20_), which is more appropriate for analysing unloaded motility data, is reported here: it is the mean of all filaments moving with a non-zero velocity after filtering out filaments with non-smooth motion (standard deviation in velocity is 20% or more than its mean velocity). When needed, myosin was deadheaded as follows: myosin was mixed with 10-20 fold molar excess of F-actin on ice for ~5 mins, followed by addition of 2 mM ATP and further 3 min incubation on ice before spinning the mixture at 95,000 rpm (Ti-100 ultra centrifugal rotor) for 20 mins at 4 °C. Unliganded myosin in the supernatant was used for motility experiments (in some cases subjected to another 1-2 rounds of deadheading if stuck percentages remained high, >10%).

### Single nucleotide turnover assay

DRX and SRX rates and proportions for WT and mutant proteins were determined from single mant-ATP turnover assays (STO), largely as described previously^38^. The SRX trends were extracted from experiments with the 15-hep constructs and the 8-hep constructs served as controls. Myosin was buffer exchanged using 100 kDa MWCO 0.5 mL Amicon filters to achieve >10^5^ dilution in STO 100 assay buffer (100 mM KOAc in 10 mM Tris-HCl pH 7.5, 4 mM MgCl_2_, 1 mM EDTA and 1 mM DTT), followed by a spin at 40,000 rpm (Ti-100 ultra centrifugal rotor) for 15 mins at 4 °C to remove protein aggregates, if any. The protein was diluted to achieve a final concentration of 500 nM in 80 μL STO assay buffer containing 6.25 mM KOAc (by mixing appropriate volumes of 0 and 100 mM KOAc containing STO assay buffers) and transferred to a well of a 96-well black plate placed in the Infinite® M Nano+ microplate fluorescence reader attached to the Te-Inject™ injector module. Software controlled injector syringes were pre-loaded with 4 μM mant-ATP ((2’-(or-3’)-*O*-(*N*-Methylanthraniloyl) Adenosine 5’-Triphosphate, Thermo-Fisher Scientific) and 40 mM ATP. Single nucleotide turnover reaction was initiated by dispensing 10 μL mant-ATP into the protein containing well from syringe A and the mixture was aged for 15 seconds before syringe B dispensed 10 μL ATP, which was immediately followed by recording fluorescence emission at 470 nm (excitation: 405 nm) every second for 16 mins. The final concentrations of different components in the reaction mixture were: 400 nM myosin, 400 nM mant-ATP, 4 mM ATP and 5 mM KOAc (or 5, 25, 100 mM in salt-dependent measurements). The injector module reduced the dead-time between adding unlabelled ATP and recording the first fluorescence reading to ~2 sec, which was manually added to the kinetic data before fitting it to an unconstrained five parameter two phase decay using GraphPad Prism. STO assays were performed with either freshly prepped protein (within 24-48 hours of protein purification) or flash frozen proteins; the data from the two was indistinguishable. 4-10 technical replicates were performed for each protein construct for each independent protein preparation (2 or more) and ambiguous fits occasionally resulting from mixing artefacts or a bubble in the optical path (< 5%) were discarded.

### Optical trapping

A dual-beam optical trap was used for harmonic force spectroscopy (HFS) measurements to obtain load dependent detachment kinetics of single myosin heads interacting with a fully stretched actin dumbbell.

Preparation of the sample chamber: First, ~1.6 μm diameter silica beads (Bangs Laboratories, Fishers, IN, USA) in a solution of 0.1% nitrocellulose and 0.1% collodion containing amyl acetate, were spin coated on top of a No. 1.5 glass coverslip. Then a sample chamber was made by sticking the silica bead coated coverslip on top of a 1 mm thick microscope slide using double-sided tape strips, while keeping the bead coated side facing the microscope slide. The tape strips were applied only on the edges of the microscope slides in order to leave enough space in the middle of the sandwich to form a chamber. Then the inside surface of the coverslip (containing the platform beads) was functionalized by flowing 15 nM SNAP-PDZ solution in AB buffer at pH 7.5 (25 mM KCl, 25 mM imidazole pH 7.5, 4 mM MgCl_2_, 1 mM EGTA, and 10 mM DTT) for ~2 minutes. The low concentration of SNAP-PDZ was used to secure only a sparse presence of the SNAP-PDZ molecules on the surface and the use of the silica beads on the cover-slip platform ensured that some of the SNAP-PDZ molecules remained elevated on top of the beads. We then flowed in 1 mg/ml BSA in AB buffer (ABBSA buffer) solution for ~5 minutes to block the remaining exposed surfaces. A 50 nM solution of WT or mutant sS1 proteins containing the C-terminal PDZ-binding domain was then flowed into the chamber for ~ 2 minutes. During this step, the sS1 molecules get translationally immobilized inside the chamber by binding to the SNAP-PDZ molecules. Then ABBSA buffer was flowed in to wash away any unbound sS1. Finally, a solution of ABBSA buffer containing streptavidin-coated polystyrene beads (1 μm diameter, Bangs Laboratories), filaments of 0.3 nM TMR-phalloidin-labelled biotinylated actin (Cytoskeleton, Denver, CO, USA), 0.018 mg/mL catalase, 0.4% glucose, 0.11 mg/mL glucose oxidase, and 2 mM ATP, was flowed in. The chamber was then sealed with vacuum grease to avoid any evaporation and taken for the HFS experiments. The entire process of chamber preparation and HFS measurements was performed at 23°C.

HFS measurements: For the HFS measurements, the two optical trap stiffnesses were kept between 0.08-0.10 pN/nm. The trap calibration was done by using the power spectrum method, described previously^52,53,55^. After the trap calibration was completed, HFS experiments were initiated by trapping a streptavidin-coated bead in each optical trap. A biotin coated actin filament was then snared between the two beads by moving the sample stage in three-dimensions, while probing the location of the filaments by phalloidin fluorescence. Then, by slowly moving the trap beads away from each other (by steering the trap-beams) the actin filament was stretched to form a “dumbbell”. While keeping the stage under 200 Hz oscillation, the actin-dumbbell was then lowered towards a platform bead (containing stationary myosin heads), to find potential myosin-actin interaction events. Upon every successful actin-myosin interaction event, brief but strong displacement would occur in both the trap bead positions simultaneously, due to their direct association with the stage oscillation. When several hundred such interactions were probed from a single myosin molecule, the time-traces of the bead-positions were saved, and the data was considered for analysis. Here we note that, to ensure that we were observing single myosin molecules interacting with any given actin dumbbell, the SNAP-PDZ concentration was kept sufficiently low that, on average, only one out of ~8-12 such trials of fishing for myosin would result in detectable and consistent interactions. The saved data (time-traces) were processed automatically, as described elsewhere^52,53,55^. Each interaction was identified by simultaneous changes in the phase and the amplitude of both the trap beads. During the period of interaction, the oscillation of the stage ensures presence of varying amounts of overall assistive or resistive forces associated with each detachment. By employing maximum likelihood estimation from the event-durations within a user-defined force bin, the rate of detachments in the presence of varied external forces were obtained. The averages of detachment rates obtained against different forces for a single molecule were then fitted with equation 1 to obtain *k*_0_ and *δ*. Further analysis of the same HFS data was done to obtain step sizes for each interacting myosin molecule. The average position and the oscillation (both amplitude and phase) of the actin-dumbbell (i.e. the actin filament stretched between two trapped beads) changes upon myosin interaction. Initially, every such myosin-actin interaction trace for one interacting myosin head were pooled together, aligned at the beginning of the binding, and extended to the length of the longest event. Then an average of the all the traces was calculated for every interacting myosin molecule. By fitting and subtracting a sine function, the oscillatory part of the average trace was then removed. The difference between the initial and final position of the sine-subtracted, averaged trace was taken as a measure of the step-size for each molecule.

### Molecular Modelling, Simulation, and Analysis

An atomic model of wild type (WT) human β-cardiac myosin in the pre-powerstroke (i.e. ADP.Pi-bound) conformation was constructed via homology modelling with *Modeller*^77^ using the 2.45 Å resolution X-ray crystal structure^47^ of pre-powerstroke bovine cardiac myosin (PDB^78^ ID: 5N69) as a modelling template. From the same X-ray structure we similarly modelled the cardiac essential light chain (ELC) and built in the disordered N-terminal extension of the ELC. Our final molecular model contains myosin residues 1-810 with ADP, H_2_PO_4_ (henceforth P_i_), and Mg^2+^ within the nucleotide binding pocket as well as residues 1-195 of the ELC. The Y115H and E497D variants were modelled using the same procedure via amino acid substitution in the modelling input sequence.

The 3 atomic models (WT, Y115H, E497D) were prepared for molecular dynamics (MD) simulation using the *tleap* module of *<u>AMBER20.</u>* Hydrogen atoms were added to the structures and then the structures were solvated in a truncated octahedral box extending at least 12 Å beyond any protein atom. Finally, counterions were added to neutralize the system. The final models contained approximately 400,000 atoms. The physical and chemical properties of the systems were described using the ff14SB^79^ (protein atoms), TIP3P^80^ (water atoms), and Li and Merz^81–83^ (ion atoms), and GAFF2 (ligand atoms) force fields. Partial atomic charges and bonded terms for ADP and P_i_ were derived from a restrained electrostatic potential (*resp*) fit to quantum mechanics calculations performed with ORCA^84^. The SHAKE algorithm was used to constrain the motion of hydrogen-containing bonds^85^. Long-range electrostatic interactions were calculated using the particle mesh Ewald (PME) method. The systems were energy-minimized, heated to 310 K in the NVT ensemble period, and density-equilibrated in the NPT as previously described. For each system, 3 500 nanosecond long production simulations were performed in the NVT ensemble using a 9 Å nonbonded cutoff and a 2 femtosecond timestep. Coordinates were saved every 1 picosecond. MD production trajectories were subsampled at 10 picosecond granularity for analysis unless specified otherwise. Interhelical angles and hydrogen bond occupancies were calculated with *cpptraj*^86^. Conformational clustering was performed using the *NMRclust* algorithm as implemented in *UCSF Chimera*^87^. Protein images were generated using *UCSF Chimera*.

### ATP binding to myosin sS1

ATP binding to WT, E497D and Y115H myosin sS1 was monitored by observing the increase in the fluorescence of mant-ATP upon binding to the myosin head. Binding experiments were performed in 10 mM Tris-HCl pH 7.5 with 4 mM MgCl_2_, 1 mM EDTA, 1 mM DTT, 100 mM KOAc. Experiments were performed on an SFM-3000 stopped-flow module (Biologic, Seyssinet-Pariset, France). Myosin sS1 (frozen or freshly prepared) was buffer exchanged into experimental buffer using 50 kDa MWCO Amicon filters (Cat # UFC5050BK, Millipore Sigma). Myosin sS1 and mant-ATP were rapidly mixed in 1: 9 or 2 : 8 ratio and injected into an observation cell coupled to a MOS-200 detection system. The final concentration of myosin in the reaction mix was 0.1 – 0.2 µM, and the reaction was carried out at 5 different concentrations of mant-ATP ranging from 0.8 - 5 µM. The mant-ATP bound to myosin head was excited by exciting the tryptophan at 290 nm which excites mant by resonance energy transfer. The fluorescence of mant was collected at 458 nm using a bandpass filter (Cat#FF01 458/64-25, Semrock). The kinetic traces were fit to a double exponential rise equation in GraphPad Prism.

## AUTHOR CONTRIBUTIONS

N.N. performed protein expression and purification, ATPase, motility and single turnover experiments, and analyzed data. D.B. performed cloning, viral production and trap experiments and analyzed data. M.C.C. performed simulations and analyzed data. R.R.G. performed stopped-flow ATP binding experiments and analyzed data. A.D. performed viral production. N.N. wrote the original draft and all authors contributed to editing and reviewing. D.B., N.N., K.M.R. and J.A.S. conceptualized and designed research and all authors contributed intellectually to discussions of the results.

## ACKNOWLEDGEMENTS

The authors thank Dr. Anne Houdusse, CNRS Research Director (CNRS/Institut Curie), for her feedback and critical inputs regarding the structural effects of the mutations, and all members of the Spudich/Ruppel lab for discussions, training and support. This work was funded by NIH grants HL117138 and 2GM033289 to J.A.S. and K.M.R.; postdoctoral fellowships from American Heart Association (Award ID 22POST908934) and Stanford MCHRI (1220552-140-DHPEU) to N.N., and a postdoctoral fellowship from American Heart Association (Award ID 23POST1027175) to R.R.G. D.B. was also funded by Rajiv Gandhi Centre for Biotechnology, Department of Biotechnology, Government of India. M. R. acknowledges funding from NIH grants GM131981 and P30 AR074990.

