## Supplemental Information for "Hypertrophic cardiomyopathy mutations Y115H and E497D disrupt the folded-back state of human β-cardiac myosin allosterically"

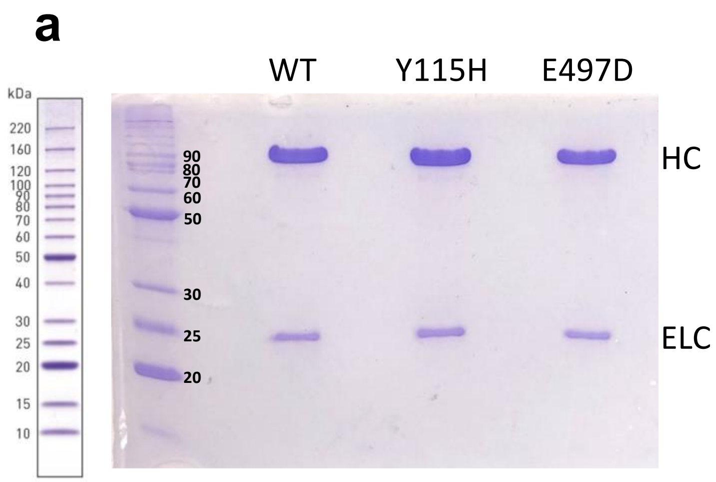


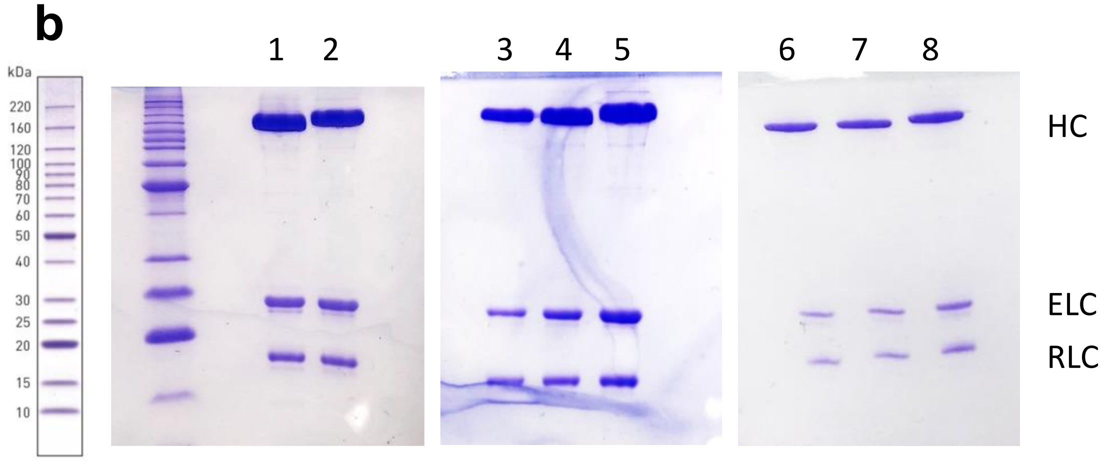


**Supplementary Figure 1. Purified sS1 and HMM constructs of myosin shown by SDS-PAGE.** (a) sS1 constructs containing β-cardiac myosin HC and ELC are shown for WT, Y115H and E497D proteins. (b) HMM constructs containing HC, ELC and RLC are shown for WT and mutants from three separate protein preparations in lanes 1-8. WT 8-hep (lane 1) and WT 15-hep (lane 2) were prepared in parallel for experiments shown in Fig. 5a, 6a and supplementary Fig. S. WT 8-hep (lane 3), Y115H 8-hep (lane 4) and Y115H 15-hep (lane 5) proteins were prepared in parallel for experiments shown in Fig. 6b. WT 8-hep (lane 6), E497D 8-hep (lane 7) and E497D 15-hep (lane 8) were prepared in parallel for experiments shown in Fig. 6c. Proteins were run on a 12.5% PAGE to visualize HC, ELC and RLC along with Invitrogen Benchmark Protein Ladder. The image on the left in (a) and (b) is a guide to compare with the molecular weight standards run on the gels.

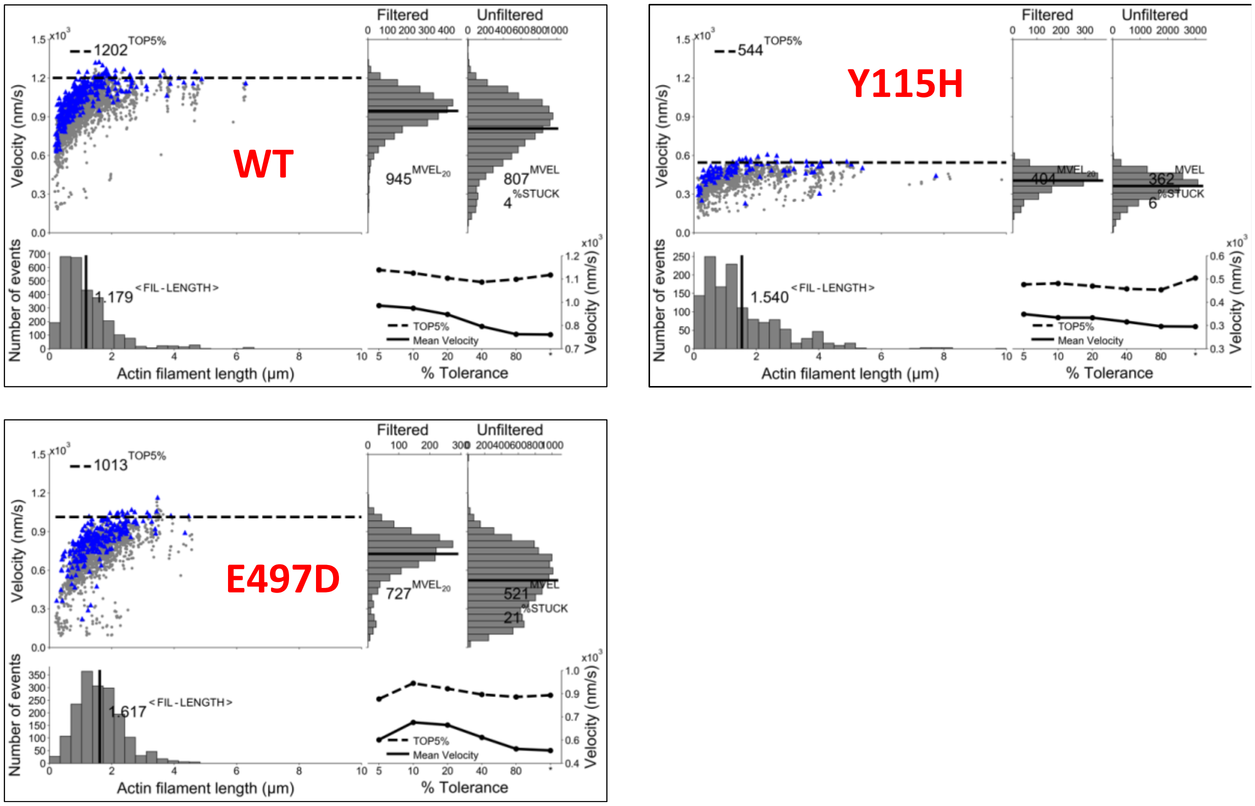


**Supplementary Figure 2. Summary of Fast Automated Spud Trekker (FAST) analysis of motility velocity data.**

FAST analysis of data from an example motility experiment movie for (a) WT, (b) Y115H, and (c) E497D sS1 proteins is shown; the movies are provided in Supplementary Movies. Parameters for FAST analysis: window size $n$ = 5, path length $p$ = 10, percent tolerance $pt$ = 20, and minimum velocity for stuck classification $minv$ = 80 nm/s. In each scatter plot on the top-left corner, $n$-frame averaged velocity value of a filament is plotted against the filament length (grey data points); each filament therefore has multiple data points depending on the number of frames in which it is seen. Blue data points represent $n$-frame averaged velocities after filtering out velocities with large fluctuations (standard deviation in velocity is 20% or more than its mean velocity when $pt$ = 20). The mean of the highest 5% values after filtering out filaments with non-smooth motion is indicated by the 'Top 5%' dashed line. The distributions of filament length, filtered velocities, and unfiltered velocities are shown, and means are indicated by the solid black line in each. %STUCK indicates the percentage of filaments whose $n$-frame averaged velocities were set to 0, which happened when velocity averaged over the entire path of a filament was lower than $minv$. These 0 velocities are not shown on the FAST plots. Bottom-right plot shows the velocity values, filtered mean velocity ($MVEL_{pt}$) and Top 5%, at different $pt$ values.

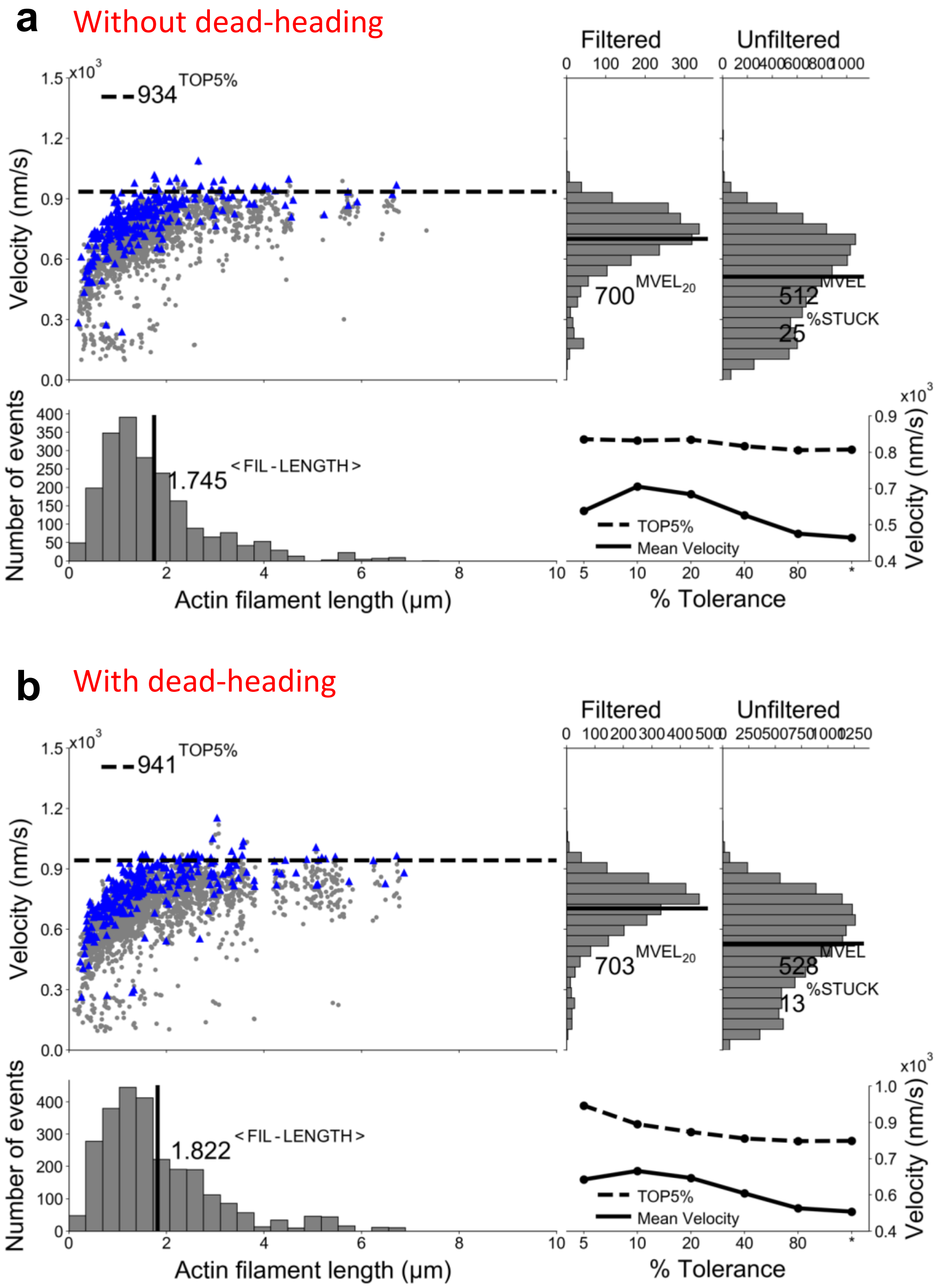


**Supplementary Figure 3. Unloaded motility velocity for E497D sS1.**
Various motility parameters from FAST analysis for E497D sS1 protein (a) without and (b) with deadheading. Here, %STUCK dropped from 26% to 13% after one round of deadheading, but no effect on any of the motility parameters was seen. See Supplementary Figure 2 legend for details.

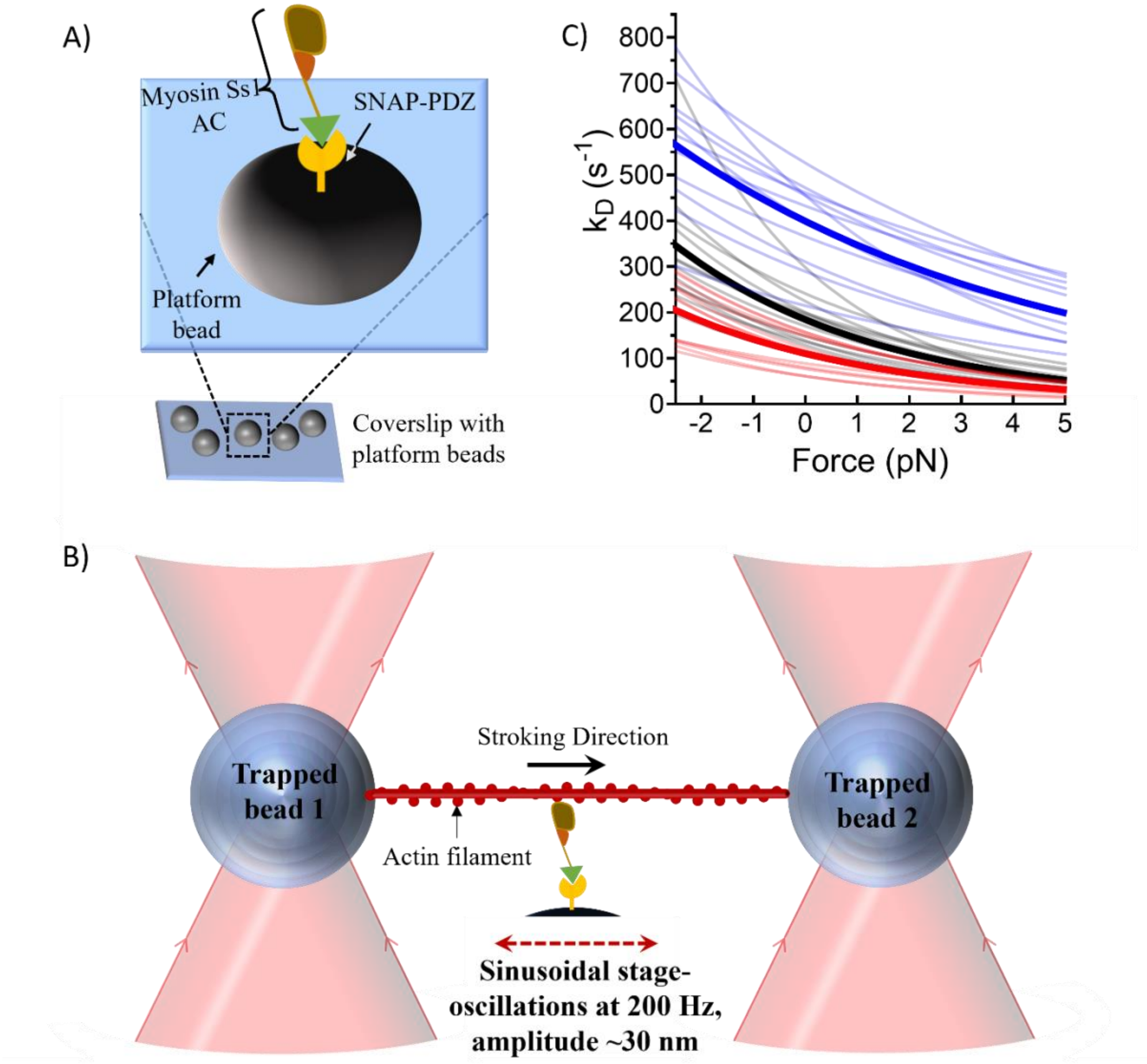


**Supplementary Figure 4. Details of the HFS setup and experiment**

(A) Orientation of a single myosin construct on top of a platform bead is depicted. The sS1 constructs have a PDZ binding domain (shown as a green triangle) that attaches with the SNAP-PDZ protein (yellow) immobilized on the surface. (B) Schematic of a typical stroking event in the HFS setup, involving a single myosin molecule, is shown. In our HFS experiment, the stage oscillates (sinusoidal) with 200 Hz frequency and with 30 nm amplitude, which results in applying a range of either assistive or resistive external forces on the stroking myosin molecules. (C) Detachment rates of the WT (grey), Y115H (red), and E497D (blue) myosin molecules is plotted in relation to the load forces. Each line in the plot results from the measurements of hundreds of stroking events from a single molecule. Average obtained from the WT, Y115H, and E497D molecules are shown as a thick black line, a thick red line, and a thick blue line, respectively.

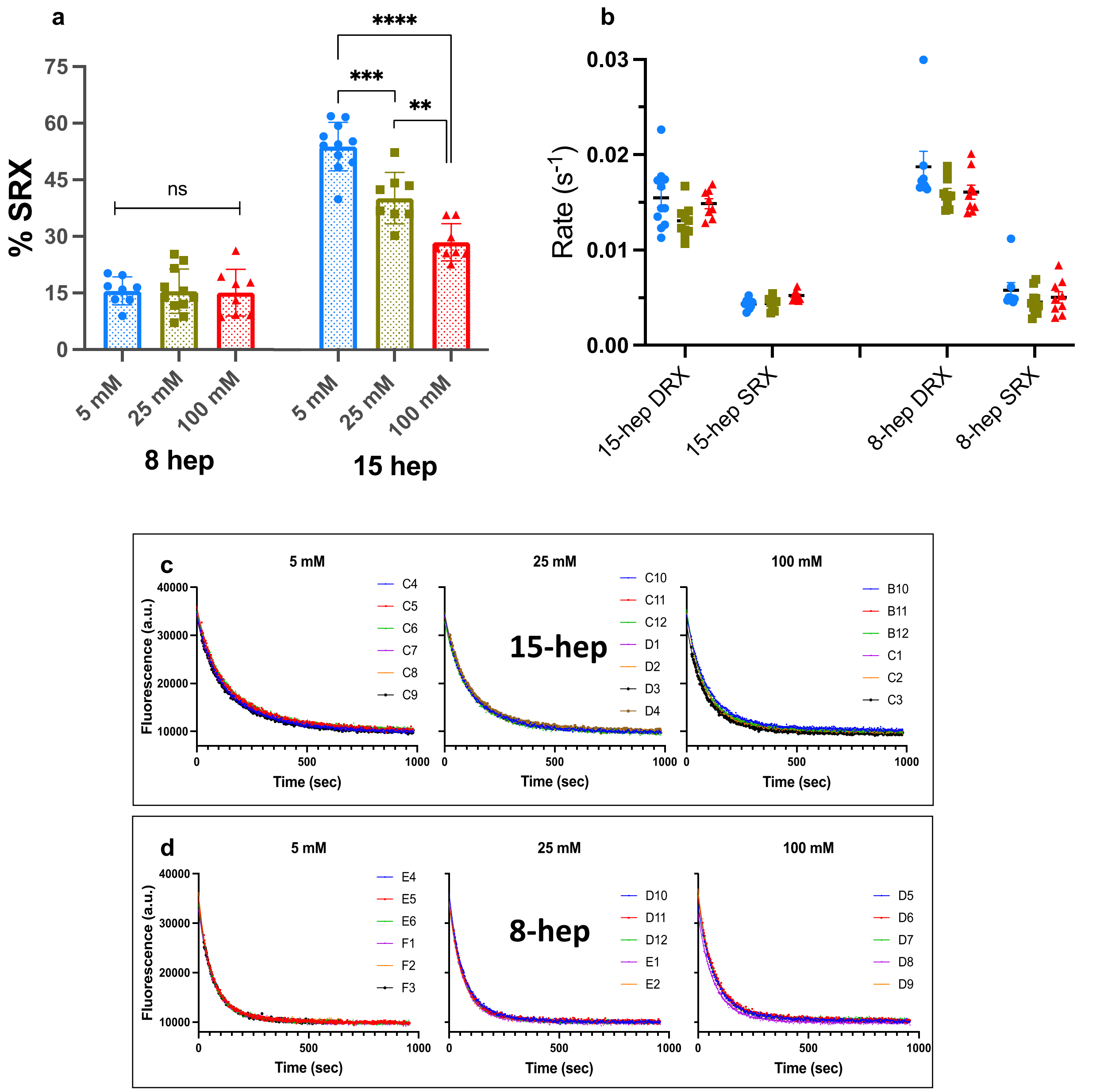


**Supplementary Figure 5. Single ATP turnover data for WT 8-hep and 15-hep proteins at different salt concentrations.**

Single turnover data was collected at 5, 25, and 100 mM potassium acetate concentrations for WT 8-hep and 15-hep proteins and the data fitted well to a biexponential decay at all salt concentrations. (a) The relative amplitude of the slow rate constants (% SRX) **and (b) DRX** and SRX rates for WT 8-hep and 15-hep proteins are shown as a function of salt. Mean ± SEM is plotted, data points represent independent measurements pooled from two independent protein preparations. ** indicates $p \leq 0.01$, *** indicates $p \leq 0.001$, **** indicates $p \leq 0.0001$. Individual raw traces at different salts fitted to biexponential decays **for (c) WT 8-hep and (d) 15-hep** from one protein preparation are shown.

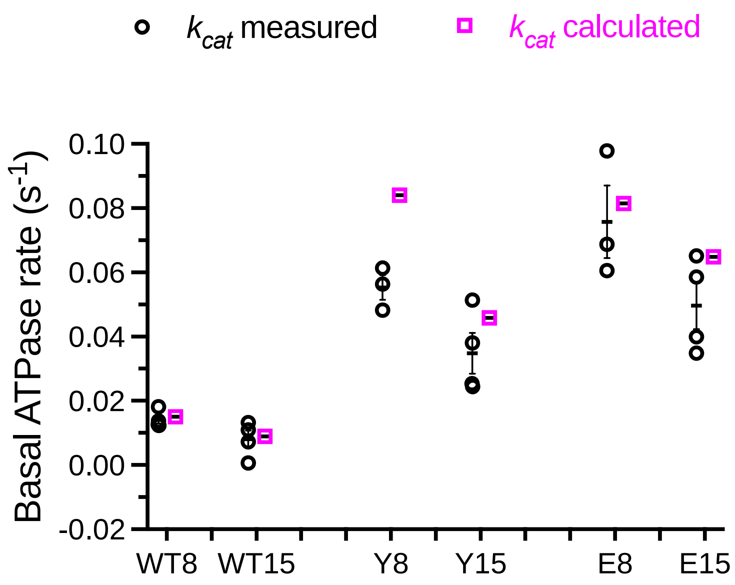


**Supplementary Figure 6. Comparison of measured basal ATPase activity and values calculated from STO measurements for HMM constructs.**

Basal ATPase rates ($k_{cat}$ measured) from the zero actin well in actin-activated ATPase measurements for WT and mutant 8-hep and 15-hep proteins (from experiments related to Fig. 6) are plotted (black circles). Each data point represents the average basal activity from three technical replicates from 1 independent protein preparation (3 or more protein preps for each protein). $k_{cat}$ calculated (pink squares) is the amplitude weighted sum of the DRX and SRX rate constants for each protein, calculated as $DRX\ fraction \times k_{fast} + SRX\ fraction \times k_{slow}$ using the single turnover data from Table 2. On the X-axis, WT8: WT 8-hep, WT15: WT 15-hep, Y8: Y115H 8-hep, Y15: Y115H 15-hep, E8: E497D 8-hep and E15: E497D 15-hep.

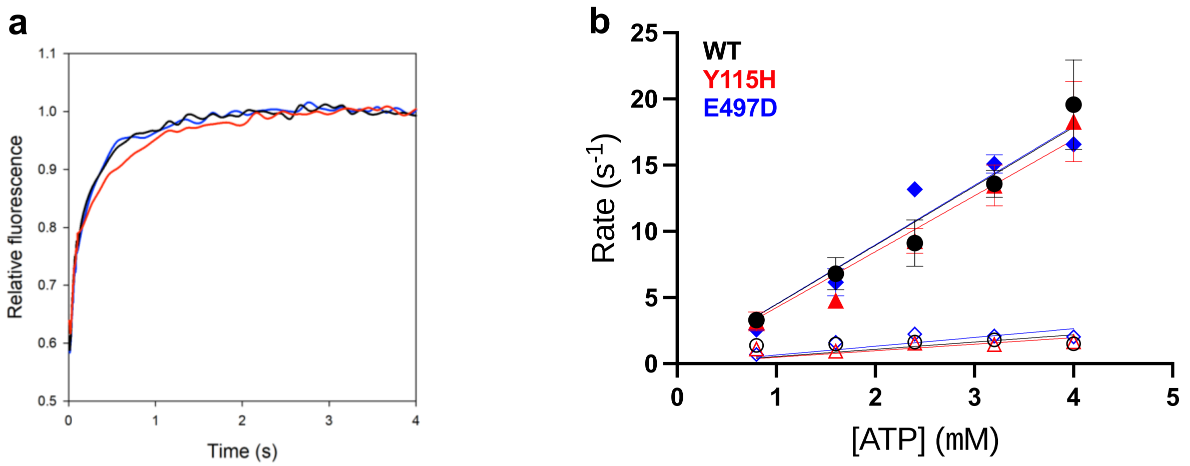


**Supplementary Figure 7. Mant-ATP binding to myosin sS1.**
ATP binding to myosin sS1 was monitored by following the increase in fluorescence of mant upon binding to the myosin head. (a) A representative kinetic trace at 3.2 μM ATP concentration is shown for WT (black), Y115H (red) and E497D (blue) sS1 proteins. The kinetic traces were fit to a double exponential equation. (b) The rates of the fast (solid symbols) and slow phases (open symbols) are shown. Mean ± SD is plotted from data collected from one protein preparation. The rate of the fast phase showed a steep dependence on ATP concentration, the data is fit to a linear regression model (passing through 0) and the slopes (representing the bimolecular rate constant) were very similar for all three proteins (WT: 0.21 $s^{-1}\mu M^{-1}$, Y115H: 0.22 $s^{-1}\mu M^{-1}$ and E497D: 0.27 $s^{-1}\mu M^{-1}$). The rate of the slow phase did not change significantly with increasing ATP concentration.

| | **WT sS1** | **Y115H sS1** | **E497D sS1** |
|---|---|---|---|
| $k_{cat}$ ($s^{-1}$) | 3.0 ± 0.18, n = 4 | ns 3.1 ± 0.11, n = 4 | ** 4.6 ± 0.28, n = 4 |
| $K_{app}$ (μM) | 19 ± 2.4, n = 4 | * 9.8 ± 0.4, n = 4 | ns 17 ± 1.3, n = 4 |
| Mean filtered velocity ($MVEL_{20}$) (nm/s) | 896 ± 23, n = 5 | **** 401 ± 10, n = 5 | *** 702 ± 28, n = 5 |
| Percent stuck filaments | < 5% | 7-10% | 12-20% |

**Table 1.** For sS1 protein constructs, $k_{cat}$ and $K_{app}$ values for WT, Y115H and E497D proteins from ATPase activity assays, each with their own independent Michaelis-Menten curve fit, from 4 independent protein preparations presented as mean ± SEM. $MVEL_{20}$ values from 5 independent experiments presented as mean ± SEM; %STUCK ranges are listed. For each mutant, whether a parameter was significantly different from WT or not is indicated: * indicates $p \leq 0.05$, ** indicates $p \leq 0.01$, *** indicates $p \leq 0.001$, **** indicates $p \leq 0.0001$, ns indicates not significantly different.

| | WT 8-hep | WT 15-hep | Y115H 8-hep | Y115H 15-hep | E497D 8-hep | E497D 15-hep |
|---|---|---|---|---|---|---|
| **$k_{fast}$ ($s^{-1}$) Fast rate, n = 5** | 0.017 ± 0.001 | 0.015 ± 0.001 | 0.09 ± 0.006 | 0.050 ± 0.003 | 0.09 ± 0.002 | 0.071 ± 0.003 |
| **$k_{slow}$ ($s^{-1}$) Slow rate, n = 5** | 0.003 ± 0.0004 | 0.004 ± 0.0001 | 0.003 ± 0.001 | 0.004 ± 0.001 | 0.004 ± 0.001 | 0.003 ± 0.0003 |
| **%SRX, n = 5** | 14 ± 2 | 55 ± 3 | 7 ± 1 | 9 ± 1 | 10 ± 2 | 9 ± 1 |
| **$k_{cat}$ ($s^{-1}$)** | 4.1 ± 0.2, n = 4 | 2.6 ± 0.1, n = 4 | 4.3 ± 0.2, n = 3 | 3.8 ± 0.3, n = 3 | 6.7 ± 0.1, n = 3 | 6.1 ± 0.3, n = 3 |
| **$K_{app}$ (µM)** | 10 ± 1, n = 4 | 14 ± 1, n = 4 | 4 ± 0.1, n = 3 | 8 ± 0.8, n = 3 | 6.1 ± 0.8, n = 3 | 11.2 ± 1, n = 3 |
| **LSAR** | | 0.63 ± 0.02, n = 4 | | 0.92 ± 0.2, n = 3 | | 0.89 ± 0.05, n = 3 |

**Table 2.** For HMM protein constructs, $k_{cat}$, $K_{app}$ and LSAR presented as mean ± SEM from *n* (indicated ) independent protein preparations, each with their own independent Michaelis-Menten curve fit. Fast rate (DRX), slow rate (SRX) and SRX fraction are presented as mean ± SEM from 5 independent protein preparations.

**Supplementary Note 1**

We note that the changes in actin-detachment kinetics and step-sizes observed in single molecule experiments account for the change observed in ensemble motility measurements for Y115H myosin, but not for E497D myosin. The presence of myosin heads in sufficient numbers on the coverslip, anchored by relatively short tethers, typically ensures that in our ensemble motility experiments the sliding velocities ($V$) of actin filaments are limited primarily by their detachment from myosin. In this experimental regime, velocity can be approximated as the step size divided by the time myosin is strongly-bound to actin ($V = d/t_s$), and since $t_s$ is inversely proportional to the detachment rate, $V = d*k_{det}$. For Y115H, a decrease in both the detachment rate as well as the step size explains the observed decrease in velocity. For E497D, an increase in ensemble velocity is expected from the increased detachment rate and unchanged step size, but the experimentally determined velocity was ~20% slower than WT myosin. The quality and smoothness of actin gliding and the concentration of myosin required to achieve that was comparable for all three proteins. The velocity was not affected by increased loading concentrations of myosin for E497D, suggesting that insufficient myosin heads on the surface is likely not the reason for the unexpectedly slow velocity. We suspect that a small fraction of E497D myosin molecules that binds irreversibly to actin in unloaded motility experiments (contributing to the increased percentage of stuck actin filaments) may act like externally added load, which coupled with the less sensitive load dependence of E497D myosin also contributes to the mismatch between the single-molecule and ensemble results.
